# Neuronal control of learned behavioral sequencing

**DOI:** 10.64898/2026.11.27.769643

**Authors:** Guillem Lopez-Fernandez, Ana Paula Menegolla, Amelie Maurel, Niniva Ghosh, Mario Martin-Fernandez, Nanci Winke, Ha-Rang Kim, Yann Humeau, Daniel Jercog, Cyril Herry, Julien Courtin

**Affiliations:** Université de Bordeaux, Neurocentre Magendie, U1215, Bordeaux, France; INSERM, Neurocentre Magendie, U1215, Bordeaux, France; Centro de Neurociencias Cajal, CSIC, Madrid, Spain; Friedrich Miescher Institute, Basel, Switzerland; Université de Bordeaux, IINS, UMR5297, Bordeaux, France; Department of Neuroscience, University of Copenhagen, Copenhagen, Denmark

## Abstract

Behavior unfolds over time as structured sequences of action continuously optimized by learning. Recent advances in behavioral segmentation tools have revealed that behavioral sequences emerge through hierarchical transitions between discrete behavioral motifs. This requires neuronal populations not only to signal and select discrete behavioral motifs, but also to prospectively coordinate transitions between them, yet the neuronal mechanisms underlying their organization remain unknown. The medial prefrontal cortex (mPFC) has been shown to track behavioral motifs and control the timing of actions, making it well suited to subserve this function. We addressed this question by combining neuronal recordings with the segmentation of behavioral sequences learned under different valence and cueing contingencies. Behavioral sequences were represented through complementary discrete and continuous coding schemes, and their structure required precise prefrontal activity patterns. Inefficient sequences were associated with stationary population dynamics, indicating a loss in the evolution of sequence representations, and optogenetic silencing of prefrontal activity impaired sequence organization, recapitulating the failure-like structure. Together, our findings uncover a general mechanism through which prefrontal neurons actively control the sequencing of learned behavior.

## INTRODUCTION

Rather than unstructured continuous motion, behavior is increasingly recognized as being composed of discrete behavioral units, often referred to as behavioral motifs, that are sequentially organized to generate a wide range of complex behaviors^1–3^. While behavioral sequences can emerge spontaneously, effective behavioral organization likely requires organisms to optimize the coordination of behavioral motifs through learning, enabling them to achieve goals, adapt to changing environments, and interact with their surroundings. These motifs are not mere kinematic patterns, but are thought to correspond to specific, internally-driven behavioral states. Behavioral segmentation tools have enabled the systematic decomposition of behavior at unprecedented resolution^4–9^, allowing the identification of specific behavioral motifs and their neuronal correlates across distributed brain regions, including the basal ganglia and cortex^10–18^.

However, adaptive behavior depends not only on the representation and selection of individual motifs, but critically relies on the neuronal mechanisms that govern transitions between them, which remain largely unknown. The medial prefrontal cortex (mPFC) is a strong candidate for this role because it tracks behavioral motifs^13–16, 18^ and exerts precise control over action timing^19–25^, suggesting that it may regulate behavioral transitions. By combining neuronal recordings in the dorsomedial PFC (dmPFC) with segmentation of behavioral responses elicited under various conditions, we identified stereotyped behavioral sequences and the neuronal computations underlying their organization. We find that prefrontal neurons represent learned behavioral sequences through complementary discrete motif representations and continuous transition coding schemes that generalize across aversive and appetitive contexts, as well as cued and self-paced regimes. Crucially, we demonstrate that prefrontal neurons prospectively control motif switches and the structuring of behavioral sequences to support adaptive performance. Specifically, disruptions in prefrontal activity, whether internally or externally generated, compromise sequence structure and lead to repetitive, inefficient behavioral patterns associated with unsuccessful outcomes. This work reveals a general mechanism by which transitions between internally-driven behavioral states are orchestrated into behavioral sequences.

## RESULTS

### Behavioral motifs organize into learned behavioral sequences

Mice were trained on a recently introduced aversive conditioning task^26^ (Figure S1A), during which animals display distinct learned behavioral sequences depending on auditory and contextual contingencies (Methods). We then applied a behavioral segmentation approach to parse mouse behavior during cued response execution (Methods), enabling the decomposition of continuous behavioral sequences into discrete behavioral motifs (Figure 1A and S1B–S1D). We identified three behavioral motifs with specific behavioral profiles (Figure 1B and S2A–S2E).

**Figure 1.**
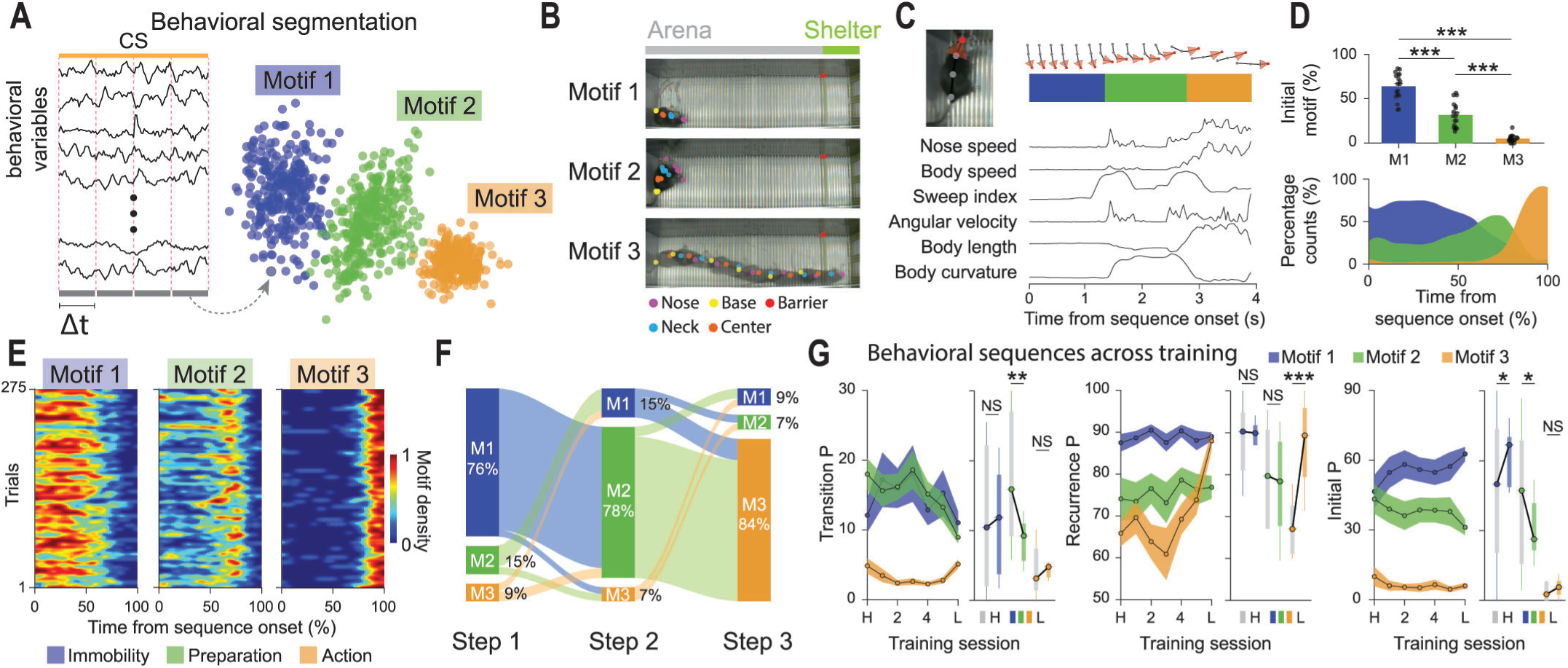
Segmentation of learned behavioral sequences. (**A**) Schematic of the behavioral segmentation pipeline. Mouse body-keypoint positions were extracted to compute behavioral variables from within-trial time-series. Data were binned (200 ms), embedded in a low-dimensional space, and clustered to identify behavioral motifs (Methods). (**B**) Example trial showing superimposed images for each motif, illustrating the corresponding behavior of the animal. Motif 1, immobility; Motif 2, preparatory head-sweeping and rotation behavior; Motif 3, action. (**C**) Example trial showing aligned body posture, behavioral motif sequence, and behavioral variables. (**D**) Top, percentage of trials initiated with Motif 1 (M1), Motif 2 (M2) or Motif 3 (M3). Circles represent individual mice. Repeated measures one-way ANOVA with post-hoc Tukey (***P<0.001). Bottom, distribution of behavioral motifs across the behavioral sequences, showing the probability of observing each motif at each time point. (**E**) Motif-density heatmaps showing the occurrence of behavioral motifs across behavioral sequences. Color intensity indicates motif density. Matrices were smoothed for visualization. (**F**) Sankey diagrams showing motif transitions across consecutive sequence positions. Columns show motif proportion at each position, and connection width indicates transition frequency. (**G**) Transition, recurrence and initial motif probabilities across training. Transition probability, switching between motifs; recurrence probability, repetition of the same motif; initial probability, occurrence at the first sequence position. For each measure, left panel shows probability from habituation to the best training day (L: learned day), and right corresponds to the quantification comparing habituation and learned day. Paired t-test or Wilcoxon signed-rank test (*P<0.05; **P<0.01; ***P<0.001; NS=not significant). Data represented as mean (bars and circles) ± S.E.M. (shaded areas and error bars). Data are from success trials; *n* = 18 mice.

Motif 1 consisted of immobility. Motif 2 was defined by preparatory actions, where mice elicited head-sweeping and body rotation movements. Motif 3 represented the output action phase of the behavioral response, characterized by high-speed directed movement. However, speed alone was insufficient to replicate the segmentation (Figure S1E), indicating that the identified motifs are not simply determined by speed. Notably, the behavioral motifs were not specific to a single behavioral context, but were consistently observed across distinct behavioral strategies (Figure S2B–S2D), suggesting that the same repertoire of motifs is employed across distinct types of responses.

Crucially, these motifs assembled into stereotyped behavioral sequences (Figure 1C–1F) from Motif 1 (immobility) to Motif 2 (preparation), and finally Motif 3 (action), suggesting a functional hierarchy in their expression (Figure 1D–1F, S2F and S2G). This sequential structure was particularly evident during active responses, while passive responses limited to immobility and preparatory behavior, without progressing to the action motif (Figure S2H). Interestingly, behavioral sequence structure was progressively shaped by learning (Figure 1G). Training reduced transitions to preparatory behavior (Motif 2), increased recurrence for the action motif (Motif 3), and shifted initial probabilities from preparatory behavior (Motif 2) to immobility (Motif 1). These learning-induced changes suggest a progressive stereotypy of learned behavioral sequences, where immobility, preparation and action follow one another with increasing efficiency through experience. Overall, these findings show that the identified behavioral motifs are not independent stereotyped behaviors, but are instead functionally organized into coherent behavioral sequences learned through training.

### Prefrontal neurons represent behavioral sequence motifs

To investigate how behavioral sequences are represented in the prefrontal cortex, we chronically implanted mice with recording electrodes in the dmPFC and analyzed neuronal activity during behavioral motif expression (Figure 2A). We observed both excitatory and inhibitory responses to behavioral motifs (Figure 2B and S3A). Notably, most neurons responded to multiple motifs, even at the single-mouse level (Figure S4A and S4B), and exhibited heterogeneous response profiles across trials (Figure S3A and S3B). Given the diversity of single-neuron responses, we applied a population decoding approach to the aggregate dmPFC neuronal activity to evaluate behavioral motif encoding (Methods). This strategy is illustrated in Figure 2D: the firing activity from the ensemble of recorded units from different mice at a given time point *t* (the so-called pseudo-population vector) is compared for different conditions, such as Motif 1 and Motif 2.

**Figure 2.**
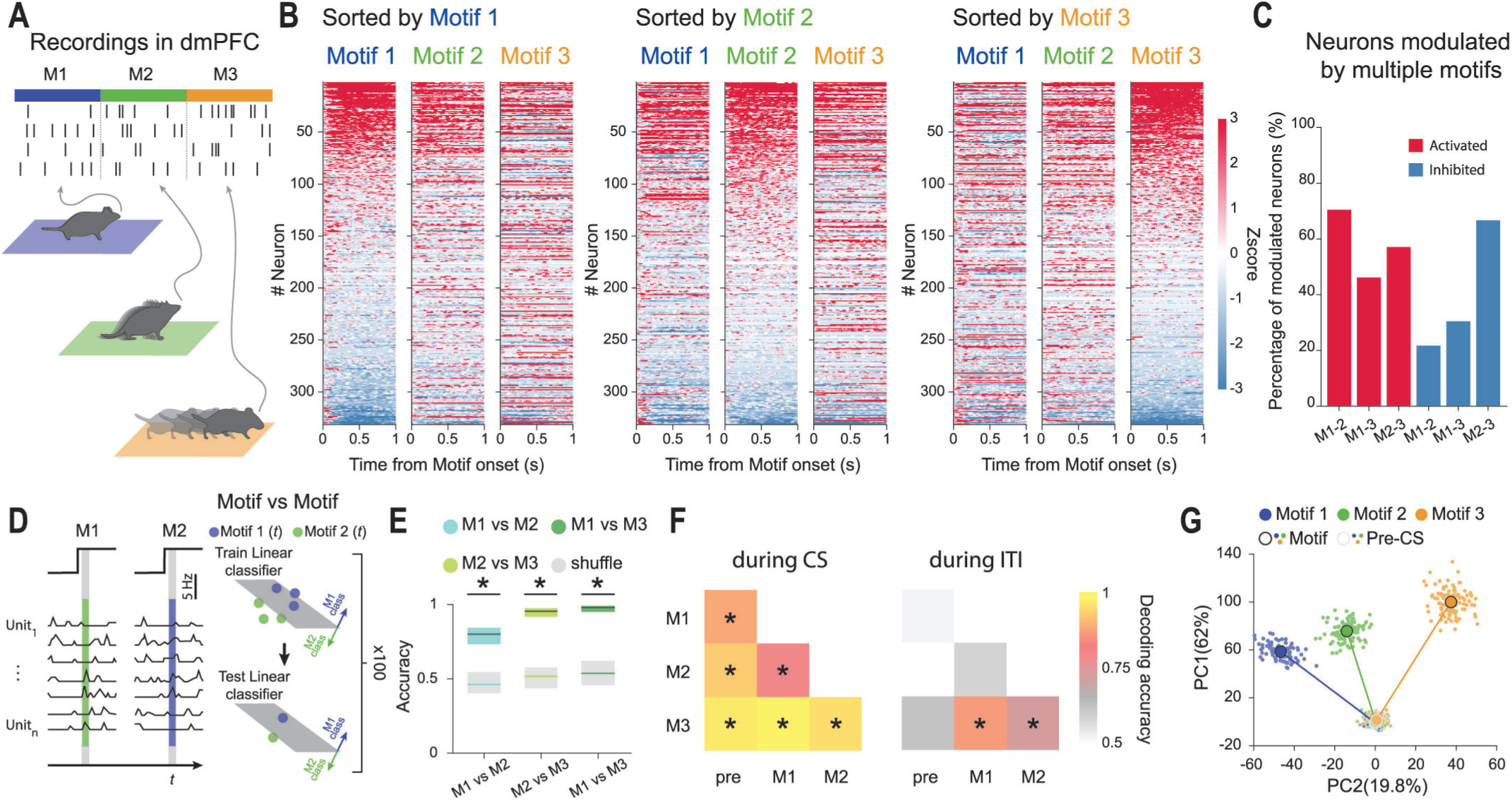
Separate representations of discrete behavioral motifs. (**A**) Schematic of dmPFC recordings aligned to motif expression. (**B**) Heatmaps of *z* scored neuronal activity aligned to motif onset. Neurons are sorted based on their activity at the onset of Motif 1 (left), Motif 2 (middle) and Motif 3 (right). Red and blue indicate increased and decreased activity, respectively. (**C**) Percentage of motif-modulated neurons responding to multiple motifs (red: activated; blue: inhibited). (**D**) Schematic of the decoding approach. The single trial-based pseudopopulation vector at time *t* is given by the spiking activity of the units for one trial. Linear classifiers are trained to classify Motif 1 and Motif 2-evoked activity (Methods). (**E**) Mean decoding accuracies for pairwise classification of behavioral motifs based on dmPFC population activity. Decoding is shown for Motif 1 versus Motif 2 (blue), Motif 2 versus Motif 3 (light green) and Motif 1 versus Motif 3 (dark green). Shuffle decoding accuracies were obtained by randomizing the condition labels (gray). Shaded areas correspond to 95% confidence intervals. (**F**) Mean decoding accuracies for motif identity during the CS period (left) and inter-trial intervals (right). Matrices show pairwise motif classification and discrimination from pre-CS baseline activity. For (E) and (F), two-sided permutation test (*P<0.05). (**G**) Projection of dmPFC population activity using PCA during pre-CS and motif-related activity in PC1 and PC2 dimensions. Large circles represent average activity, small circles represent resampled population vectors. Data are from success trials; *n* = 331 neurons from 14 mice.

Using this approach, we were able to decode all three motifs from one another (Figure 2E and 2F) and from pre-CS baseline activity (Figure 2F, S3C and S3D) with high decoding accuracy, suggesting distinct representations for each behavioral motif. Importantly, decoding remained robust at the single-mouse level (Figure 4C–4F). In contrast, decoding of motifs outside CS presentation was not achievable except for the high-speed Motif 3 (Figure 2F, S3D and S3F), suggesting a state-dependent encoding of motif-related activity. Notably, removing speed-modulated neurons reduced baseline decoding of Motif 3 to chance but spared CS-period decoding (Figure S3G and S3H), indicating that motif representations during trial execution reflect genuine differences in motif-specific activity beyond locomotion. Importantly, motif representations generalized across distinct behavioral strategies (Figure S3E), including both active and passive responses. To further evaluate whether prefrontal neurons distinctly represent behavioral motifs, we applied principal component analysis^27^ (PCA; Methods) to reduce the dimensionality of instantaneous pseudo-population vectors during motif expression (Figure 2G). The resulting PCs (Figure S3I) consistently separated motif responses from pre-CS activity and from one another (Figure 2G, S3J and S3K). Together, these findings suggest that behavioral motifs within learned sequences reflect specific behavioral states that are separately represented in the dmPFC.

### Population activity predicts behavioral transitions

Given that prefrontal activity encodes individual behavioral motifs, we next asked whether these representations contribute to the organization of behavior across time. Specifically, we examined whether prefrontal activity predicts behavioral transitions, thereby implicating prefrontal networks in generating structured behavioral sequences rather than passively reflecting ongoing actions. We first observed a gradual recruitment of activated and inhibited neurons during transitions (Figure 3A and S5A). Unlike CS onset-evoked activity, where most neuronal responses were locked to CS presentation (Figure 3B), the majority of transition-modulated neurons responded prior to the execution of the upcoming motif (Figure 3A, 3B and S5A), suggesting a prospective shift in population activity. To test this possibility, we used a population decoding approach to evaluate motif encoding during behavioral transitions. By training a decoder to classify each 200-ms time bin relative to the preceding bin, we tracked the evolving structure of population activity over time (Figure 3C). This revealed significant decoding across successive time bins leading up to behavioral transitions, indicating that population activity gradually evolves to represent the upcoming motif (Figure 3D). In contrast, decoding from pre-CS to Motif 1 expression increased sharply at CS onset, consistent with the external sensory cue driving the neuronal response (Figure 3D). These results suggest that dmPFC population activity predicts upcoming behavioral elements through internally generated, anticipatory dynamics. To further test this, we performed cross-temporal decoding of behavioral transitions. We sequentially trained a decoder to classify each time bin (100 ms) from a motif expressed during the transition, and tested each classifier with the remaining time bins across the same transition (Figure 3E). Accordingly, dmPFC activity progressively shifted away from preceding motif activity while simultaneously aligning with upcoming motif representations (Figure 3E, 3F, S5B and S5C). This indicates that, instead of simply anticipating movement, dmPFC activity is continually reorganized in relation to the current and upcoming behavioral states. This pattern was observed in both active and passive responses, indicating a common mechanism across behavioral strategies (Figure S5D–S5F). Contrasting with this, cross-temporal decoding before motif transitions remained stable over time (Figure S5G–S5I), indicating that these gradual changes are specific to the transition period. Together, these results reveal a parallel progression of dmPFC representations from past to future behavioral states preceding motif transitions, supporting a role for prefrontal neurons in actively structuring motif switches to form learned sequences.

**Figure 3.**
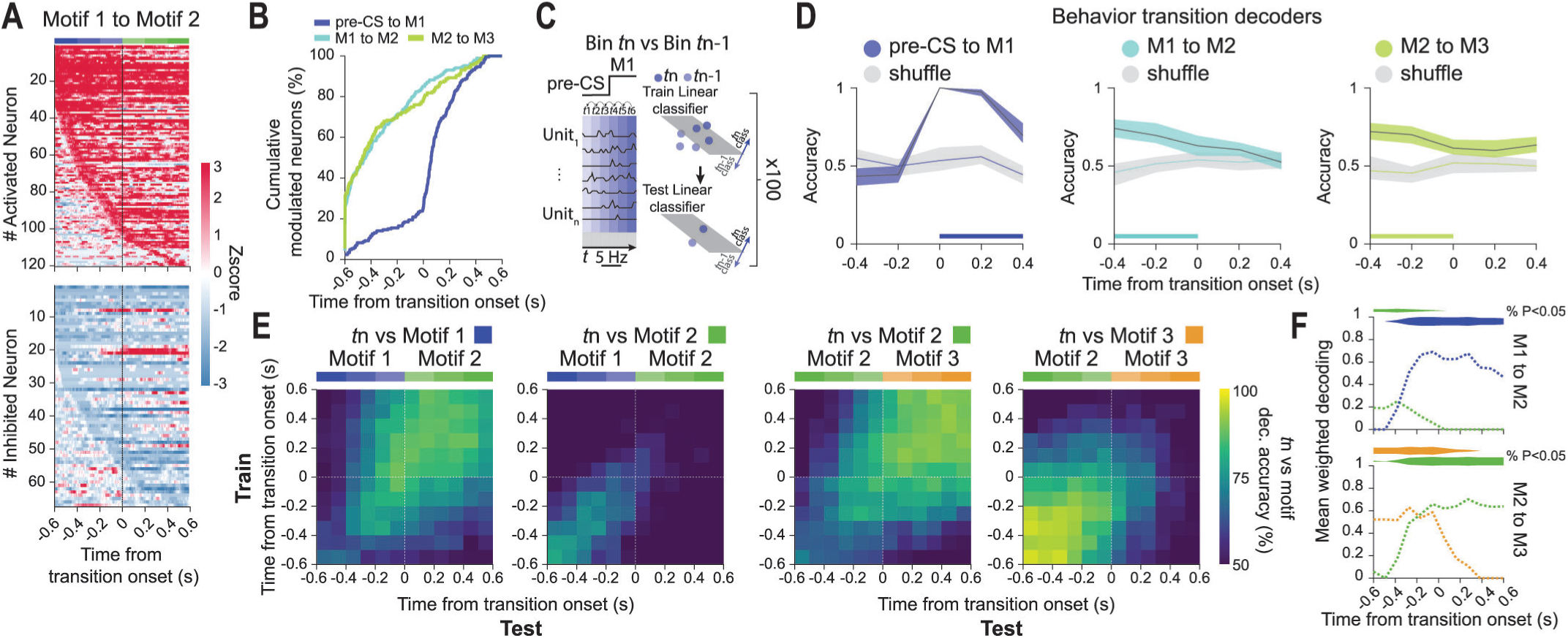
Anticipatory representations of behavioral transitions. (**A**) Heatmaps of neuronal activity for activated (top) and inhibited (bottom) neurons, aligned to transitions from Motif 1 to Motif 2. Neurons are sorted by the first detected modulation time. Red and blue indicate increased and decreased activity, respectively. (**B**) Cumulative percentage of dmPFC-modulated neurons during periods from pre-CS to Motif 1 (dark blue), Motif 1 to Motif 2 (light blue) and Motif 2 to Motif 3 (green), aligned to transition onset. (**C**) Schematic of the decoding approach. The single-trial pseudopopulation vector at time *t* is given by the spiking activity of the units for one trial during a given behavioral transition. Linear classifiers are trained to classify each time bin *t*n (200-ms bin) of the transition from the preceding bin *t*n-1 (Methods), enabling examination of the time-to-time evolution of population activity during the transition period. The example illustrates decoding of Motif 1 to Motif 2 transitions. (**D**) Mean decoding accuracies for consecutive time bins during behavioral transitions. Decoding is shown for pre-CS to Motif 1 (dark blue), Motif 1 to Motif 2 (light blue) and Motif 2 to Motif 3 (green) transitions. Shuffle decoding accuracies were obtained by randomizing the condition labels (light gray). Shaded areas correspond to 95% confidence intervals. Horizontal lines indicate significant decoding accuracies (two-sided permutation test; P<0.05). (**E**) Cross-temporal decoding of behavioral transitions (100-ms bin) from motif population activity for Motif 1 to Motif 2 (left) and Motif 2 to Motif 3 transitions (right). Classifiers were trained at one time point and tested across all others for each transition bin (Methods). (**F**) Quantification of cross-temporal decoding over time for transitions from Motif 1 to Motif 2 (top) and Motif 2 to Motif 3 (bottom). Mean weighted decoding accuracy was calculated for each test time bin as the average decoding accuracy across significant bins in the corresponding column, weighted by the proportion of significant bins. Horizontal shaded areas indicate significant decoding, and height of shaded area indicates the percentage of decoded bins (two-sided permutation test; P<0.05). Data shown are from success trials; *n* = 318 neurons from 13 mice.

### Behavioral sequence structure predicts trial outcome

To test the functional relevance of these findings, we compared behavioral sequences across success and failure outcomes, where failure was defined as the selection of an erroneous response. Failure trials exhibited key differences in behavioral sequence structure (Figure 4A and 4B), with increased preparatory behavior (Motif 2) and reduced immobility (Motif 1) and action (Motif 3). Moreover, we observed a disruption in the stereotyped arrangement of these motifs, with a marked reduction in the proportion of sequences progressing from immobility to preparation to action (Figure 4C). Accordingly, transition and recurrence probabilities for Motif 2 increased in failure sequences (Figure 4D), suggesting facilitated transitions into preparatory behavior and reduced transitions out of it. To confirm this, we quantified the number of transitions and time spent in Motif 2 across mice. Both metrics increased in failure relative to success sequences (Figure 4E), and these effects remained consistent across active and passive responses (Figure S6G–S6I). This suggests a bias toward entering and remaining in preparatory behaviors during unsuccessful performance. Notably, Motif 2 expression in success trials also increased after extinction (Figure 4F), suggesting that sequence structure is modulated by task contingencies. To further assess the specificity of behavioral sequences to trial outcome, we implemented a leave-one-out classification approach across mice to predict trial outcome from the observed sequences. Models trained on all but one mouse were used to predict sequences from the held-out mouse as success or failure, yielding an estimate of outcome prediction accuracy (Methods). This approach accurately predicted trial outcome (Figure 4G and S6J), indicating that sequence structure reliably distinguishes success from failure trials. Together, these results indicate that successful performance relies on specific behavioral sequences shaped by learning and maintained under reinforcement, whereas deviations in sequence structure results in suboptimal behavior.

**Figure 4.**
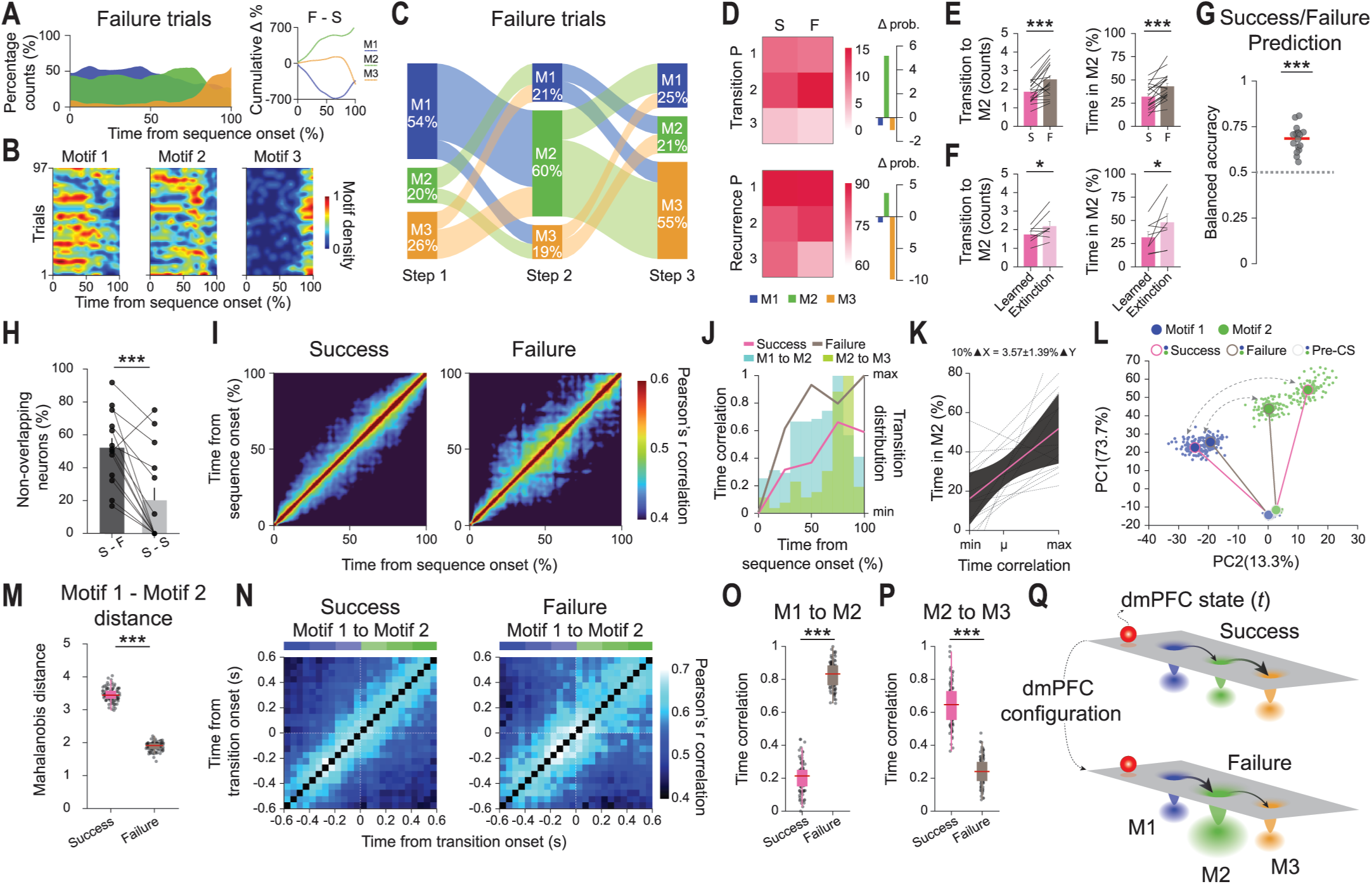
Behavioral sequence structure and neuronal activity differ across trial outcomes. (**A**) Left, distribution of behavioral motifs across failure behavioral sequences. Right, cumulative difference in motif probability between success and failure trials. (**B**) Motif-density heatmaps during failure trials. Color intensity indicates motif density. Matrices were smoothed for visualization. (**C**) Sankey diagrams of motif transitions during failure trials. Columns show motif proportion at each position, and connection height indicates transition frequency. (**D**) Transition (top) and recurrence (bottom) probability heatmaps across success and failure trials (left), and their differences (right). (**E**) Number of transitions to Motif 2 (left) and time expressing Motif 2 (right) during success (pink) and failure (brown) trials. Bars represent mean ± S.E.M., lines indicate individual mice. Paired t-test (***P<0.001). (**F**) As in (E), but comparing learned (pink) and extinction (light pink) in success trials. Paired t-test (*P<0.05). (**G**) Trial outcome prediction from behavioral sequences (Methods). Circles indicate held-out mice and red line indicates mean balanced accuracy across mice. One-sample t-test against chance (0.5; ***P<0.001). For (A–E) and (G), *n* = 18 mice. For (F), *n* = 8 mice. (**H**) Percentage of non-overlapping transition-modulated neurons across outcomes versus a within-condition control. Non-overlapping neurons were modulated in one condition but not the other. Bars represent mean, circles represent individual mice. (**I**) Mean correlation matrices of population activity during success (left) and failure (right) sequences. (**J**) Time correlation during behavioral sequences with the distribution of Motif 1 to Motif 2 (blue) and Motif 2 to Motif 3 transitions (green). (**K**) Relationship between dmPFC time correlation (mean-centered; μ) and Motif 2 expression during success trials estimated with a linear mixed-effects model (Methods). Black lines indicate individual mouse fits, pink line indicates the population estimate, and shaded area represents the 95% confidence interval. A 10% increase in time correlation predicted a 3.57 ± 1.39% increase in Motif 2 expression (LME: β = 49 ± 19.1, t = 2.57, P = 0.01). (**L**) Projection of dmPFC population activity using PCA during pre-CS and motif-related activity in success and failure trials. Large circles represent average activity, small circles represent resampled population vectors. (**M**) Mahalanobis distance between Motif 1 and Motif 2 population vectors. Box plot center line represents the median and box limits are 25th and 75th percentiles. (**N**) Mean correlation matrices of single-trial pseudo-population vectors during Motif 1 to Motif 2 transitions in success (left) and failure (right) trials. (**O**) Time correlation during Motif 1 to Motif 2 transitions in success and failure trials. (**P**) As in (O), but for Motif 2 to Motif 3 transitions. For (H, M, O and P), paired t-test or Wilcoxon signed-rank test (***P<0.001). For (H–P), *n* = 331 neurons from 14 mice. (**Q**) Conceptual model of dmPFC population dynamics during behavioral sequences. The schematic depicts the instantaneous dmPFC population activity (red sphere) moving through an activity landscape. Valleys correspond to behavioral motifs’ associated activity patterns. In success trials, valleys have similar widths and depths, generating similarly stable behavioral states. In failure trials, Motif 2 valley is deeper and broader, promoting transitions into and persistence within this state.

To control for potential segmentation bias arising from fewer failure trials (Figure S6A), failure motifs were assigned using k-Nearest Neighbors (k-NN) interpolation onto the success-derived behavioral feature space (Methods). Importantly, motif labels remained consistent when segmentation was performed using all trials (Figure S6B–S6D).

### Prefrontal neurons orchestrate behavioral sequences

Notably, although behavioral sequences differed by outcome, motif behavioral profiles remained consistent (Figure S6E and S6F), suggesting that motifs capture similar underlying behavioral states independent of trial outcome. Consistent with this, motif-related population activity during failure trials strongly generalized to decoders trained on success trials (Figure S7A), indicating that despite differing in their sequential organization, success and failure trials share the same constituent states.

These observations thus indicate that failure does not emerge from the absence of appropriate behavioral states, but from differences in how they are implemented and coordinated over time. This predicts that altered dmPFC population dynamics might drive the unstructured behavioral sequences associated with failure trials. Aligned with this hypothesis, many transition-modulated neurons in success trials were unresponsive during failure (Figure 4H, S7B and S7C). This occurred despite similar overall neuronal recruitment patterns across outcomes (Figure S7F and S7G), indicating a partial remapping of neuronal ensembles while preserving population-level dynamics during motif transitions. To further investigate this, we quantified the temporal structure of dmPFC activity by computing pairwise Pearson correlations between population vectors across trial time points (‘time correlation’; Figure 4I). In this framework, higher correlations between distant time points indicate more stationary activity patterns, whereas lower correlations reflect a more dynamic progression through population states. Notably, failure trials exhibited higher correlations, indicating more stationary activity, whereas success trials showed faster decorrelation of activity, consistent with a stronger progression of dmPFC activity patterns over time (Figure 4I and 4J). Moreover, across success trials, higher time correlations predicted longer Motif 2 occupancy (Figure 4K, S7D and S7E), linking stationary prefrontal activity to persistence in preparatory behavior. Consistent with this, PCA revealed a reduced distance between Motif 1 and Motif 2 population activity during failure (Figure 4L and 4M), indicating that transitions between these states require smaller changes in population activity. These data suggest that stationary prefrontal activity, characteristic of failure trials, is associated with prolonged occupancy of preparatory states and may contribute to the repetitive behavioral sequences observed in these conditions.

Interestingly, differences in time correlation between outcomes were largest when most transitions from Motif 1 to Motif 2 occurred and minimal during Motif 2 to Motif 3 transitions (Figure 4J), suggesting transition-specific differences in population activity. To investigate this, we constructed pseudo-population vectors across mice and examined population dynamics during specific motif transitions (Figure 4N). Consistent with our previous observations, failure trials exhibited higher time correlations during Motif 1 to Motif 2 transitions, indicating more stationary population dynamics and a stronger convergence toward the Motif 2 state (Figure 4N and 4O). In contrast, transitions from Motif 2 to Motif 3 showed the opposite pattern (Figure 4P), suggesting a reduced convergence onto the output Motif 3 and an increased persistence of the Motif 2 state in failure trials. Together, these findings suggest that failures are driven by a distinct dmPFC configuration that stabilizes preparatory behavior (Figure 4Q), promoting entry into while limiting exit from this behavioral state, and thereby contributing to the unstructured and repetitive behavioral sequences observed during unsuccessful responses. In contrast, successful performance requires a dynamic reconfiguration of dmPFC activity patterns as the behavioral sequence unfolds.

### Prefrontal activity disruption induces failure-like behavioral sequences

To test the causal role of prefrontal neurons in orchestrating behavioral sequences, we optogenetically silenced dmPFC activity using the inhibitory opsin ArchT. Crucially, dmPFC inhibition disrupted success behavioral sequences, replicating the behavioral structure of failure trials by increasing both the transition and recurrence probabilities of Motif 2 (Figure 5A and S8A). Furthermore, dmPFC silencing reduced Motif 1 while increasing Motif 2 expression compared to non-stimulated trials and GFP controls (Figure 5B–5D and S8C–S8F). Importantly, dmPFC inhibition increased the number of transitions and time spent in Motif 2 (Figure 5E), with no effect in GFP controls (Figure 5F) or failure trials (Figure S8B). Moreover, light stimulation increased Motif 2 expression to failure-trial levels, such that no significant differences were observed between light-on success and light-off failure trials (Figure 5E).

**Figure 5.**
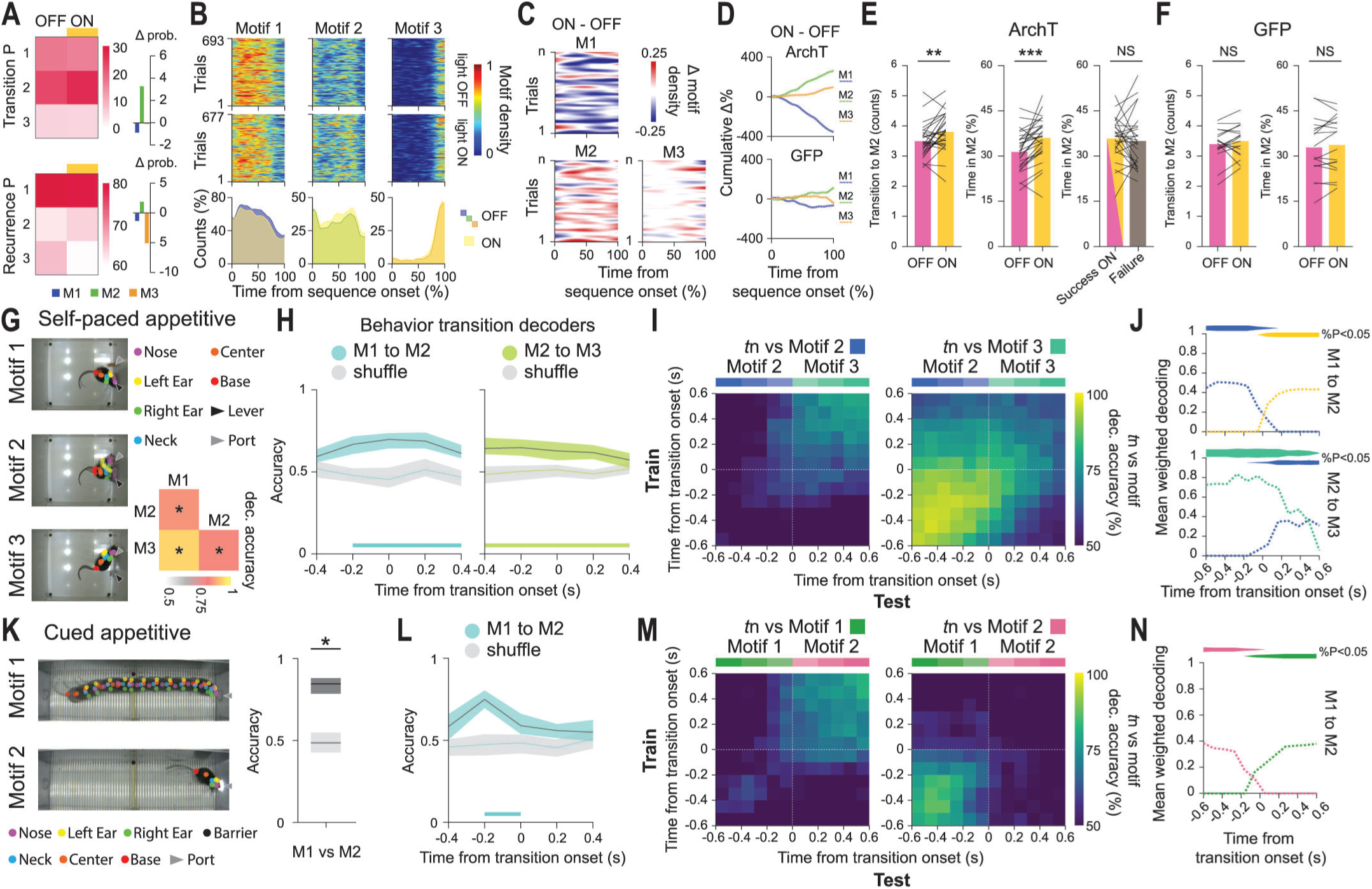
Optogenetic disruption of behavioral sequence structure and generalization of behavioral transition representations. (**A**) Left, heatmaps of transition (top) and recurrence (bottom) probabilities for behavioral motifs in light-off and light-on conditions during success trials. Right, difference in motif probabilities between stimulation conditions. (**B**) Top, motif-density heatmaps showing the occurrence of behavioral motifs across behavioral sequences in light-off and light-on conditions. Color intensity indicates motif density. Matrices were smoothed for visualization. Bottom, distribution of behavioral motifs over time in light-off (blue: Motif 1; green: Motif 2; orange: Motif 3) and light-on (yellow) conditions. (**C**) Difference heatmaps between light-on and light-off conditions. Red and blue indicate increased and decreased motif density during inhibition, respectively. Matrices were smoothed for visualization. (**D**) Cumulative difference in motif probability between light-off and light-on trials for ArchT (top) and GFP (bottom) groups. (**E**) Left, number of transitions to Motif 2 during success light-off (pink) and light-on (yellow) trials in the ArchT group. Middle, percentage of time expressing Motif 2 during success light-off and light-on trials. Right, percentage of time expressing Motif 2 during success light-on (pink-yellow) and failure light-off (brown) trials. Bars represent mean ± S.E.M, lines indicate individual mice. Paired t-test (**P<0.01; ***P<0.001; NS=not significant). (**F**) As in (E), but for GFP group. Paired t-test (NS=not significant). For (A–F), *n* = 29 ArchT and 18 GFP mice. (**G**) Left, example trial showing superimposed images for behavioral motifs identified in a self-paced appetitive task. Motif 1, lever-pressing; Motif 2, turning; Motif 3, consumption. Right, decoding accuracies for motif identity from pairwise classifications. (**H**) Decoding accuracies across consecutive time bins (200-ms bin) during Motif 1 to Motif 2 (blue) and Motif 2 to Motif 3 transitions (green). Shuffle decoding was obtained by randomizing condition labels (light gray). Shaded areas correspond to 95% confidence intervals. Horizontal lines indicate significant decoding (two-sided permutation test; P<0.05). (**I**) Cross-temporal decoding (100-ms bin) for transitions from Motif 2 to Motif 3. Classifiers were trained at one time point and tested across all others for each transition bin (Methods). (**J**) Quantification of cross-temporal decoding for Motif 1 to Motif 2 (top) and Motif 2 to Motif 3 transitions (bottom). Mean weighted decoding accuracy was calculated for each test time bin as the average decoding accuracy across significant bins in the corresponding column, weighted by the proportion of significant bins. Horizontal shaded areas indicate significant decoding, and height of shaded area indicates the percentage of decoded bins (two-sided permutation test; P<0.05); *n* = 10 mice and 1776 neurons. (**K–N**), As in (G–J), but for a cued appetitive task. Motif 1, reward approach; Motif 2, consumption. Two-sided permutation test (P<0.05); *n* = 4 mice and 349 neurons.

Together, these data demonstrate a causal involvement of dmPFC neurons in coordinating behavioral sequences for the adaptive execution of behavior. Disruptions in dmPFC activity, whether internally (during failures) or externally generated (during manipulation), result in repetitive behavioral patterns that impair motif sequence structure.

### Prefrontal control of learned behavioral sequences to self-paced and cued reward

Having established a role for the dmPFC in controlling behavioral transitions in aversive contexts, we next asked whether this function generalizes to other learned behavioral sequences and appetitive contexts. For this, we parsed behavioral sequences elicited within two reward-guided paradigms and examined the underlying dmPFC activity. In a self-paced appetitive task^28^, mice autonomously obtained rewards by pressing a lever and subsequently visiting a reward port, with no explicit stimuli signaling trial start or reward availability. Mice stereotypically expressed behavioral sequences consisting of lever press (Motif 1), turning (Motif 2), and consuming the reward (Motif 3; Figure 5G, S9A, S9B and S9E). In a cued appetitive task^29^, mice in a shuttle box responded to an auditory conditioned stimulus associated with a reward, where shuttling to the opposite compartment led to reward availability. Behavioral sequences primarily consisted of reward approach and consumption, which were most often expressed in stereotyped succession (Figure 5K, S9C, S9D and S9F). Consistent with our observations in the aversive task, all motifs across both tasks could be decoded from one another with high decoding accuracy, indicating separate dmPFC representations (Figure 5G and 5K). Using the same transition-decoding approach as in the aversive task (Figure 3C), we observed significant pre-transition decoding in both reward-based sequences (Figure 5H and 5L), indicating that dmPFC activity progressively evolved toward the upcoming motif before its execution. Moreover, cross-temporal decoding revealed reciprocal changes in motif decoding, indicating gradual shifts in dmPFC representations from the preceding to the upcoming motif (Figure 5I, 5J, 5M, 5N and S9G). By contrast, in a contextual fear conditioning task, prefrontal activity was tightly locked to freezing onset and offset, lacking the anticipatory and progressive decoding dynamics observed for learned behavioral sequences (Figure S9H–S9M). This suggests that prefrontal activity preferentially controls transitions within learned, rather than innate behavioral sequences.

Overall, these results extend our findings from aversive to reward cued and self-paced behavioral sequences, supporting a common prefrontal mechanism for orchestrating learned behavioral structure.

## DISCUSSION

Here, by combining behavioral sequence segmentation with neuronal recordings, we demonstrate that the prefrontal cortex represents not only discrete behavioral motifs but also their transitions in a progressive and anticipatory fashion, and that dmPFC activity actively contributes to the coordination of these motifs into structured behavioral sequences. By replicating these findings across aversive and appetitive contexts, as well as cued and uncued regimes, our data support a general mechanism through which prefrontal neurons flexibly orchestrate the sequencing of learned behavior.

At the behavioral level, our results indicate that diverse behavioral outputs can be decomposed into a limited set of reusable behavioral motifs. We identified a common repertoire of behavioral motifs across distinct behavioral strategies and outcomes, highlighting that complex behavior arises from the flexible organization of a shared repertoire of discrete actions^4^. Crucially, we found that learning shapes the structure of behavioral sequences. Training progressively biased motif transitions toward successful sequences, whereas extinction restored an early-training, failure-like sequence structure. These findings suggest that optimal performance depends not only on learning task contingencies and outcomes, but on learning to efficiently structure a conserved motif repertoire into specific behavioral sequences.

Our observation that dmPFC neurons respond to multiple behavioral motifs is aligned with previous reports of overlapping motif-tuned ensembles in the dmPFC^13^. Despite this overlap, behavioral motifs remained encoded by separate representations at the population level. By providing evidence for prefrontal motif representations across distinct behavioral sequences spanning positive and negative valence, as well as cued and uncued conditions, our findings extend previous work linking neuronal activity to the expression of behavioral motifs in various brain regions^10–18^. The absence of these representations during baseline conditions argues against a purely kinematic-based interpretation and suggests that they emerge when motifs are expressed within learned behavioral sequences. Accordingly, motif encoding persisted after removing speed-modulated neurons, indicating that the identified behavioral states are independent of speed.

Contrasting with cue-triggered responses, in which most neurons were modulated at CS onset, behavioral transitions showed gradual neuronal recruitment over time. Within this organization, dmPFC neurons responded in anticipation to motif expression, and upcoming motifs could be decoded before their execution, indicating that prefrontal activity predicts upcoming motif transitions rather than merely reflecting ongoing behavior. This anticipatory coding evolved progressively and was confined to the transition period, with representations of the current motif gradually decreasing as representations of the upcoming motif increased. Moreover, neuronal activity differed across transition types, suggesting that dmPFC neurons not only gate behavioral transitions, but also shape them through transition-specific activity patterns. These findings suggest that, although individual behavioral motifs are represented as discrete states, motifs are linked through a continuous transition process, during which representations of past and future behavioral states simultaneously evolve in opposite directions. Previous work reported changes in dmPFC activity before running during classical active avoidance responses^30^, but did not examine prefrontal activity according to individual behavioral states or their temporal evolution. By explicitly parsing behavior, our results demonstrate that dmPFC activity dynamically evolves in relation to specific behavioral states within learned sequences, rather than simply preceding movement. Crucially, we replicated these findings across behavioral sequences elicited under aversive, appetitive, cued and uncued conditions, providing evidence for a general mechanism underlying the organization of behavioral sequences. However, this pattern was absent for freezing transitions. Although freezing could be decoded with high accuracy, the anticipatory and progressive evolution of state representations observed for learned behavioral sequences were largely lacking. This distinction suggests that the dmPFC preferentially supports the organization of learned behavioral sequences, where flexible coordination between discrete motifs is required, while transitions between innate states may be signaled without being actively orchestrated through the same organizing dynamics. This is consistent with recent findings showing that, during spontaneous behavior, dmPFC behavioral state-tuned activity emerges only after the onset of behavioral transitions^18^. In contrast, learning to produce purposeful behavioral sequences, which requires prospective behavioral planning, appears to recruit the prefrontal cortex in a fundamentally different way. Thus, these data suggest that prefrontal neurons actively participate in the sequencing of learned behavior.

Supporting this interpretation, failure trials were associated with disruptions in motif transitions at both the behavioral and neuronal level. Moreover, optogenetic inhibition of dmPFC activity disrupted motif transitions and generated failure-like behavioral sequences. In essence, failure sequences were characterized by the recurrent expression of preparatory behavior (Motif 2), indicative of a less efficient behavioral structure that resembled that observed in early training. This was accompanied by altered dmPFC population dynamics, suggesting that suboptimal prefrontal activity results in repetitive behavioral patterns that impair motif sequence structure. These observations highlight a possible link between prefrontal activity stationarity and behavioral rigidity, a finding with potential clinical implications, and suggest that unsuccessful performance is implemented by altered prefrontal dynamics (Fig. 4Q). Aligned with these observations, optogenetic dmPFC silencing decreased task performance^26^ and disrupted behavioral sequence structure, mirroring the altered sequence dynamics observed during failure trials. Together, these data reveal an active control of prefrontal neurons in coordinating behavioral sequence transitions to support adaptive performance.

An important consideration of our study, and for behavioral segmentation approaches more generally, is that behavioral motifs can arbitrarily be subdivided into increasingly finer-grained components, raising the possibility that identified motifs comprise multiple sub-states. As segmentation becomes more granular, however, motifs may increasingly capture kinematic features rather than meaningful behavioral representations. Here, we focused on motifs reflecting broad behavioral states to investigate sequence organization, but future studies should determine how motif granularity relates to internal states across multiple scales (but see Weinreb et al.^18^).

Behavioral segmentation has substantially advanced the study of spontaneous behavior^10–12, 18^, but has been applied less frequently to controlled behavioral paradigms, where neuronal activity is typically analyzed in relation to the onset and offset of stimulus-evoked events. Our data, however, indicate that animals transition through various behavioral states, representing a potential source of unaccounted variability. Thus, incorporating behavioral segmentation into the routine of systems neuroscience might be a crucial step toward better disentangling variables of interest from changes in behavioral state. Moreover, unlike paradigms in which action sequences are prescribed by the experimental design, the sequences identified here emerged spontaneously from the animals’ behavioral strategy, thus reflecting self-organized behavioral structure rather than being externally dictated. Despite other studies have shown neuronal correlates of behavioral states and sequences^13, 14, 18, 28, 31, 32^, we here provide a direct link between learned behavioral structure and the neuronal computations underlying its organization. Collectively, our data establish a general mechanism through which the prefrontal cortex prospectively coordinates internally-driven motifs of action into structured behavioral sequences, generating both discrete and continuous sequence representations required for flexibly orchestrating learned behavior.

## ACKNOWLEDGMENTS

We thank all members of the C. Herry laboratory for comments and helpful discussions. We thank S. Laumond, J.Tessaire, C.Dupuy and all the technical staff of the housing and experimental animal facility of the Neurocentre Magendie.

## AUTHOR CONTRIBUTIONS

Conceptualization: G.L.F., C.H., J.C. Methodology: G.L.F., M.M.F., Y.H., D.J., J.C. Investigation: A.M., N.G., A.P.M., N.W., H.R.K., G.L.F. Formal analysis: G.L.F. Software: G.L.F. Supervision: C.H., J.C. Writing – original draft: G.L.F., C.H., J.C.

## METHODS

All procedures were performed in accordance with standard ethical guidelines (European Communities Directive 86/60-EEC) and were approved by the committee on Animal Health and Care of the Institut National de la Santé et de la Recherche Médicale and the French Ministry of Agriculture and Forestry (agreement no. A3312001).

### Animals

Mice trained in the aversive conditioning task (multi input-output (MIO) paradigm; ^26^) were male C57BL6/J mice (8–12 weeks of age, Janvier) individually housed under a 12-h light–dark cycle and provided with food and water ad libitum. The housing temperature was 22 ± 1 °C, and housing humidity was 60% ± 5%. For the experiments and subsequent behavioral segmentation, we used 18 mice, of which 14 were implanted for electrophysiology recordings. An additional mouse implanted with a gradient-index (GRIN) lens for calcium imaging recordings was used for single-mouse neuronal analyses. For optogenetic experiments, we used 29 mice injected with ArchT, and 18 control mice injected with GFP. Mice trained in the contextual fear conditioning task and cued appetitive task were 19 and 4 male C57BL6/J mice (Janvier), respectively, implanted for electrophysiology recordings. Mice trained in the self-paced appetitive task were 10 male C57BL6/J mice (Envigo) implanted with GRIN lenses for calcium imaging recordings.

### Behavior

Mice were trained in an aversive conditioning task (MIO paradigm) in which they learned to select the correct behavioral response by integrating context and tone contingencies. The MIO paradigm was performed in a behavioral box (40 × 10 × 30 cm) composed of Plexiglas walls and a grid floor used to deliver foot shocks (0.6 mA). The box was divided into an arena zone (35 × 10 cm) and a shelter zone (5 × 10 cm) by a small plastic hurdle (0.5 cm high). The shelter zone was marked with visual cues to facilitate its recognition. The box was equipped with infrared beams to detect mice shuttling between the shelter and arena zones. The behavioral box was enclosed in an acoustic isolated box containing speakers, a light source and a video camera (30 frames per second; Cineplex, Plexon) above the behavioral box. Three different auditory cues were delivered at a volume of 75 dB: CS^+^1 (7 kHz), CS^+^2 (2 kHz) and CS^−^ (white noise). The auditory cues were presented for 7 s or until mice shuttled from one zone to the other (shelter and arena) in the case of the CS^+^. The CS^−^ was presented for 7 s independently of mouse behavior.

The presence of the auditory cue and the location of mice in either the shelter or the arena established the different trial types. In active trials, a CS^+^1 was presented when mice were in the arena. Mice had to shuttle to the shelter before the end of the CS (7 s) to avoid a foot shock (1 s). In passive trials, the CS^+^1 was presented when mice were in the shelter. Mice had to remain in the shelter for the duration of the CS (7 s) to avoid a foot shock (1 s). In inverse trials, a CS^+^2 tone was presented when mice were in the shelter. Mice had to shuttle to the arena before the end of the CS (7 s) to avoid a foot shock (1 s). In control trials, the CS^−^ was presented either in the shelter or in the arena without any reinforcement and independently of mouse behavior. The foot shocks lasted for a maximum of 1 s (shuttling to the correct area of the box would end the foot shock) and co-terminated with the CS when mice did not perform the correct behavioral response for each CS^+^ trial type (failure trials). Therefore, at the end of failure trials, there was an extra second of CS^+^–US co-presented. The different trial types were intermingled and dependent on mouse location and tone presentations. During each behavior training session, mice were exposed to 90 intermingled and pseudorandomized ‘tone-presentation events’ of three different types: 30 CS^+^1 presentations: active or passive trials depending on whether mouse was in the arena or shelter, respectively; 30 CS^+^1 presentation if mice were in the arena and CS^+^2 if mice were in the shelter: active or inverse trials, respectively; and 30 control CS^−^ presentation independent of mouse location. The ITI was pseudorandomized, ranging from 35 to 80 s. A behavioral session therefore lasted approximately 75 min on average. The sequence of tone presentations and ITIs was altered across training to prevent the learning of a trial type sequence. The training sessions consisted of a habituation session (H; identical to a normal training session but lacking shock presentations) and seven to eight training sessions. A group of mice was then submitted to extinction training consisting of three to four sessions identical to normal training sessions without shock presentation. The experimental box was cleaned with 70% ethanol before and after each session.

To assess the generalization of dmPFC behavioral sequence representations, we analyzed neuronal activity recorded during three additional behavioral paradigms: contextual fear conditioning, a self-paced appetitive task, and a cued appetitive task. For contextual fear conditioning, mice were exposed to a conditioning context in which they received aversive foot shocks and were subsequently re-exposed to the same context to assess conditioned freezing responses. The context consisted of an open-ceiling cube (28 x 28 x 50 cm; length x width x height), with a grid floor connected to a shocker (Coulbourn Instruments) for the delivery of mild electric foot shocks (0.7 mA, 1 s). During the conditioning session, mice received five foot shocks distributed over a 9 min period. The following day, mice underwent a retrieval session in which they were re-exposed to the same context for 3 min, and freezing behavior was assessed.

For the self-paced appetitive task^28^, mice were trained in an apparatus containing a lever and a lick port on the same wall to autonomously obtain rewards in a goal-directed manner. Mice were food restricted to 85% of their free-feeding body weight 4 days before and throughout the behavioral experiments and fed ∼2 hours after their daily behavioral sessions with ∼2.5 g of regular food. Training consisted of constant reinforcement in which a sucrose reward (20 μl, at 20%) was delivered after lever pressing. Behavior was entirely self-paced, without explicit stimuli signaling trial start or reward availability. For the cued appetitive task, mice were trained in an instrumental approach and avoidance task^29^. Mice first underwent differential avoidance conditioning (CS^+/–^). Subsequently, mice were water-restricted for a maximum of 15 consecutive days, with body weight maintained at ≥ 85% of baseline. Mice then underwent an approach conditioning procedure in which a CSr (13kHz) signaled water reward availability (80 μl) upon shuttling to the opposite compartment. A nose poke on the port of the shuttled compartment before 12 seconds from the onset of CSr triggered the delivery of the water reward. After reaching stable CS shuttling probability for the CSr (80%), mice were submitted to an avoidance-approach session consisting of a block of randomized CS^+^ and CS^−^ presentations (25 trials each), followed by a block of CSr and CS^−^ presentations (25 trials each), as well as a session consisting of all four CS types (50 trials each) presented intermingled. Subsequent analyses were performed using pooled data from both sessions.

### Pose estimation and behavioral segmentation

Behavioral tracking was performed using DeepLabCut^33^ to track the mice’s movements during the training sessions. We selected four body keypoints for tracking and subsequent behavioral segmentation: the mouse’s nose, neck, center and base of the tail. An additional keypoint marked the position of the barrier separating the arena from the shelter compartment of the training box. This yielded the estimated position in X and Y directions for each of the selected keypoints in each frame. To control for keypoint detection errors, we treated trackpoints for which the predicted position differed by more than 30 pixels from the previous frame as missing points, and estimated their position with linear interpolation between the last known position and next. The pose estimation of each mouse and session was then subsampled to 5 Hz (200-ms bins) for subsequent behavioral segmentation. To perform behavioral segmentation, we selected the day for each mouse on which performance was highest across the three types of threatening trials (L: learned day), ensuring there were enough samples of each behavioral response—approximately a minimum of 200 samples per response—for subsequent low-dimensional embedding and clustering. Next, we measured a total of thirty behavioral variables from the estimated keypoint positions: absolute speed in X and Y direction, distance between body markers, cumulative position in X and Y direction, cumulative distance between body markers, average angle between body markers and cumulative average angle between body markers (Figure S2A). We used winsorization to limit the influence of outliers and min-max normalization to scale all behavioral variables between 0 and 1. We performed behavioral segmentation in each mouse separately on time-series restricted to active, passive, inverse success and control trials lasting a minimum of 3 s (CS onset–CS offset). For trials in which mice exited the shelter (inverse success trials), the analysis window extended by 1 s beyond CS offset to ensure that the complete behavioral sequence, including Motif 3, was captured. Because this additional period would overlap with shock delivery in passive failure trials, all analyses involving failure trials were restricted to the interval between trial onset and offset (CS onset to before US onset), thereby avoiding contamination by shock. To avoid biasing our segmentation towards control (CS^−^) trials, we randomly selected 10 control trials (350 samples) to perform the segmentation. Next, to visualize and segment mouse behavior, we applied t-distributed Stochastic Neighbor Embedding (t-SNE) to the 30-dimensional behavioral variable space. t-SNE was implemented using the following parameters: perplexity = 60, learning rate = 1000, number of dimensions = 3, and distance = correlation. The resulting 3D embedding preserved local behavioral similarities and enabled qualitative assessment and clustering of stereotyped behavioral motifs. Following t-SNE embedding, we applied agglomerative hierarchical clustering on the 3D coordinates to identify discrete groups of similar behavioral patterns with a fixed number of 8 clusters based on empirical tuning. These clusters captured local neighborhoods within the behavioral embedding while avoiding excessive fragmentation, and served as an initial unsupervised segmentation. To incorporate prior knowledge of behavioral categories of interest (‘immobility’ and ‘action’), we performed a semi-supervised consolidation of the initial clusters. Clusters that were predominantly composed of ‘immobility’ samples were merged into a single immobility cluster (Motif 1). Clusters that were predominantly composed of ‘action’ samples were merged into a single action cluster (Motif 3). The remaining clusters located between Cluster 1 and Cluster 3 were merged into a single cluster that reflected a combined behavioral profile related to head sweep and turning behavior (preparatory behavior, Motif 2). This resulted in a final set of three clusters (behavioral motifs). This segmentation yielded a sequence of discrete behavioral motifs for each trial, representing the evolving structure of behavioral responses. Cluster histograms were performed by computing Principal Component Analysis (PCA) on the centroids of the three identified clusters and then projecting the 3D coordinates obtained with t-SNE onto the first principal component of the centroids, representing the best projection axis. To control that the identified behavioral motifs were not only dependent on speed differences we applied the same protocol as described above using only behavioral variables related to speed (speed in X and Y directions; 8 variables).

For the segmentation of behavioral responses during the self-paced and cued appetitive tasks, we applied the same approach as described above but using an extra pair of keypoints including left and right ears. Behavioral variables were adapted to the behavioral characteristics of each task but using a similar set of variables to those used in the aversive task. For the self-paced task, trials were defined as the period spanning the first lever press to the offset of reward availability. For the cued appetitive task, trials were defined from CS onset to one second after consumption onset, and consumption was defined a posteriori as the period spanning one second after the first lick post shuttling.

For identification of the behavioral motifs during failure trials, ITI periods and the rest of training and extinction sessions, we followed the same procedure as described above but, for each mouse, we used k-Nearest Neighbors (k-NN) to interpolate the multi-behavioral variable samples onto the low dimensional space previously obtained with t-SNE in the learned day. k-NN was implemented with 5 neighbors and using Euclidean distance as the measure of similarity between points. Neighbor contributions were weighted by their distance, assigning greater influence to closer neighbors. To control that this procedure yielded similar results as using t-SNE, we repeated the same behavioral segmentation procedure using balanced success, failure and control trials on the best day of performance and applied t-SNE to obtain a lower-dimensional map of all the samples (Figure S6A–S6D). The percentage of match between the two methods was calculated as the percentage of failure samples assigned to the same behavioral motif using either k-NN or t-SNE. We quantified a similarity index between success and failure behavioral motifs by computing, for each mouse, the mean absolute difference between success and failure behavioral variable values minus one.

For identification of behavioral motifs in mice submitted to optogenetic experiments, we followed the same procedure as described above, but using reference motif centroids obtained from the non-optogenetic dataset for each trial type. Each high-dimensional behavioral sample was assigned to a reference motif using a weighted majority vote from k-NN (k = 3) distances, in which neighbors contributed according to the inverse of their Euclidean distance. To assess whether a trial was sufficiently represented by the reference dataset, we calculated the mean Euclidean distance to neighbors belonging to the assigned motif, normalized by the median distance among reference samples of the assigned motif, yielding a relative distance that accounts for differences in motif dispersion. The median normalized distance across all samples within a trial was then computed. Trials whose median normalized distance exceeded the median reference normalized distance by more than two standard deviations were excluded from the analysis. This procedure yielded a common behavioral motif labeling across light-on and light-off conditions, enabling unbiased comparison of sequence structure between lighting conditions.

### Behavioral analyses

We calculated the performance as the number of correct trials over the total number of trials for each trial type. The global performance was quantified as the average performance of the different threatening trial types (active, passive and inverse trials). To analyze the shuttling of mice during control trials (during the CS^−^ presentation), we considered that they shuttled when they changed the area where they were located (shelter or arena) during the trial. Immobility and action measurements were computed from the filtered coordinates of the mouse keypoint body markers (nose, neck, center and base of the mouse). For immobility analysis, mice were considered to be immobile if the mean speed of their nose and center was below 1 cm/s for at least 1 second. For action analysis, action initiation was detected in active responses (active success and inverse success) if the speed of the mouse center was above a 95% confidence interval yielded by the speed over the duration of each trial, and the mouse crossed the barrier during the period of speed increase. The action period was considered from the detected moment of action initiation until trial offset.

For freezing detection in mice submitted to contextual fear conditioning, instantaneous speed (pixels/frame) was computed from the X-Y coordinates of the nose and center smoothed over 5 consecutive frames. Freezing was defined as periods during which center speed was <0.65 pixels/frame and nose speed simultaneously <0.5 pixels/frame, for a minimum duration of 1 s.

These speed thresholds were set empirically, and were further controlled on live video recordings. The combined nose and center criterion excluded immobility periods unrelated to freezing, such as grooming or sniffing, and animals were considered freezing only when displaying complete body immobility except for respiratory movements.

To obtain transition, recurrence and initial probabilities, a transition count matrix was constructed for each behavioral sequence by counting every transition between consecutive motifs. Counts were summed across all sequences of a given trial type and normalized by the total number of transitions originating from each motif to obtain the transition probability matrix. For each motif, the transition probability was quantified as the average probability of transitioning into that motif from any other motif across consecutive steps (i.e., the average of the corresponding column of the transition matrix, excluding the diagonal). Recurrence probability was defined as the probability of remaining in the same motif across consecutive steps (i.e., the diagonal elements of the transition matrix). Initial probability was calculated as the fraction of sequences that started with each motif. Thus, transition, recurrence and initial probabilities respectively quantified the likelihood of transitioning into a motif, remaining in the same motif, and occupying the first position in a sequence. Global probabilities were calculated by averaging across trial types (active, passive and inverse trials).

Sankey diagrams were used to illustrate behavioral sequence structure. For each trial, we extracted three-step sequences in which the three motifs were expressed in succession. Then, we calculated motif proportions at each step (represented by the column height) and the percentage of transitions between motifs from each step to the next (represented by the connection height). To visualize the distribution of behavioral motifs over time, individual behavioral sequences were time-normalized to a fixed length of 70 samples by resampling each sequence at uniformly spaced positions and assigning each new position the value of the nearest original sample. For each motif, binary matrices indicating motif presence over time were generated across trials and smoothed with a 2-D Gaussian smoothing kernel to improve visualization of motif occurrence across sequences. The resulting smoothed matrices were displayed as heatmaps, with color intensity indicating motif density. The number of transitions to Motif 2 was defined as the number of distinct entries into Motif 2 averaged across trials for each mouse. The percentage of time expressing Motif 2 was quantified as the proportion of Motif 2 in each behavioral sequence averaged across trials for each mouse. Global metrics for success and failure trials were obtained by averaging across trial types (active, passive and inverse trials).

To characterize motif-related behavior, we measured distinct behavioral variables aligned to the expression of the identified behavioral motifs. Instantaneous nose and body speeds were depicted as the median speed across all instances of each motif. Sweep index was calculated from nose lateral motion relative to the body axis. The resulting lateral displacement was smoothed and quantified over a sliding window as the range between maximum and minimum lateral positions (sweep amplitude). Sweep amplitude was then normalized by a forward-motion factor to reduce the influence of forward locomotion onto lateral movement detection. The final sweep index was calculated as a bounded index between 0 and 1 using an exponential transformation, with higher values indicating stronger lateral sweeping movements. Angular velocity was calculated as the absolute rate of change in the animal’s body orientation angle over time, with orientation defined by the neck-to-center body axis. Body rotation was quantified from the animal’s body orientation angle, calculated from the neck-to-center body axis, providing a measure of the animal’s rotational posture. Body length was quantified as the distance from nose to base of the mouse.

Body curvature was quantified as the angle between the front and rear body axes. The front and rear body vectors were calculated from neck to nose and from base to center, respectively, and the angle between them resulted from their dot product. Behavioral variables were subsequently quantified by averaging their values over two periods relative to motif expression: the pre-motif period was defined as the 1 s interval preceding motif onset, whereas the motif period corresponded to the entire motif expression period.

### Hidden Markov Models

To model the transition dynamics underlying observed behavioral sequences, we employed Hidden Markov models (HMMs). Each model assumes that the observed behavioral sequence is generated by an underlying Markov process transitioning among a set of unobserved latent states, each associated with a probability distribution over the observed behavioral motifs.

Separate HMMs were fit for each trial type (active, passive and inverse trials for success and failure outcomes), allowing us to characterize how behavioral structure differs across success and failure sequences. Model fitting was performed using MATLAB’s built-in function hmmtrain (Mathworks). This function applies the Baum-Welch algorithm to estimate the state transition matrix, which represents the probability of transitioning from one latent sate to another, and the emission matrix, representing the probability of observing each behavioral motif given a latent sate. The state transition probability matrix was then used to calculate the models’ transition and recurrence probabilities for the different behavioral motifs. Comparisons between success and failure trials were made between trial types yielding the most similar behavioral output (active success versus active failure; inverse success versus passive failure; passive success versus inverse failure).

To further investigate the structure and dynamics captured by the fitted HMMs, we generated synthetic Markov sequences for each trial type. Motif 1 was specified as the initial state because it was the most frequently observed initial motif (Figure 1D). We simulated sequences with a fixed duration of 7 s using the state transition and emission matrices estimated by the HMM from each trial type, thereby generating synthetic sequences representative of each fitted model. Each synthetic sequence was produced according to the estimated transition matrix, and at each step a behavioral motif was sampled from the emission distribution associated with the current latent state. These model-generated sequences were used to evaluate whether the models could reproduce key features of the observed behavioral sequences.

To assess the trial outcome predictive power of the obtained behavioral sequences, we implemented a leave-one-out cross-validation approach at the subject level. In each iteration, one mouse was held out as the test set, while the remaining mice formed the training set. For each trial type, an HMM was trained using all trials from the training set as described above. Then, to predict the outcome of the test mouse’s behavioral sequences, we used the MATLAB’s built-in function hmmdecode (Mathworks) to compute the log-likelihood of an observed sequence given a trained HMM. Each test sequence was evaluated using both the success-trained and failure-trained HMMs corresponding to the same trial type comparison (active success versus active failure; inverse success versus passive failure; passive success versus inverse failure). The predicted label (success or failure) was assigned based on which model yielded the higher likelihood for the observed sequence. Prediction accuracy was quantified separately for each held-out mouse. Balanced accuracy was calculated as the average of the accuracies for correctly classifying success and failure trials, thereby giving equal weight to both outcome classes despite different number of trials. Mouse-level balanced accuracies were compared against chance (0.5) using a two-sided one-sample t-test.

### Surgeries and recordings for in vivo electrophysiology

Mice were anesthetized with isoflurane (induction, 3%; maintenance, 1.75%) in O2. Body temperature was maintained at 37 °C with a Temperature Controller System (FHC), and eyes were hydrated with Ocry-gel (TVM). Mice were placed in a stereotaxic frame (Kopf Instruments), and three stainless-steel screws were attached to the skull. Following craniotomy, mice were bilaterally or unilaterally implanted in the left dmPFC with an electrode array at the following coordinates relative to the bregma: +2.15 mm AP, ±0.7 mm ML and −1.25 mm DV from the dura with an angle of 14°. Each electrode array consisted of 16 individually insulated nichrome wires (13 μm in diameter, impedance of 30–100 kΩ, Kanthal) fixed to an electrode guide attached to an 18-pin connector (Omnetics). Connectors were referenced or grounded via a silver wire (127 μm in diameter, A-M Systems) placed above the cerebellum. All implants were secured using Super-Bond cement (Sun Medical). During surgery, long-and short-lasting analgesic agents were injected (Metacam, Boehringer; Lurocaïne, Vetoquinol). After surgery, mice were allowed to recover for at least 7 d. Electrodes were connected to a headstage (Plexon) containing 16 unity-gain operational amplifiers. Each headstage was connected to a 16-channel PBX preamplifier (gain 1,000x, Plexon) with bandpass filters at 300 Hz and 8 kHz. Spiking activity was digitized at 40 kHz and isolated by time–amplitude window discrimination and template matching using an OmniPlex system (Plexon). At the conclusion of the experiment, electrolytic lesions were induced before transcardial perfusion to verify electrode tip location using standard histological techniques.

### Virus injections and optogenetics

For optical silencing of dmPFC CaMKIIa-expressing neurons, we bilaterally injected 0.15–0.2 μL of either GFP (AAV5-CaMKIIa-GFP, titer of 1.5 × 10^12^, UNC Vector Core Facility) or ArchT (AAV5-CaMKIIa-ArchT-GFP, titer of 5.3 × 1012, Addgene) into the dmPFC (relative to the bregma, +2.15 mm AP, ±0.7 mm ML and −1.3 mm DV from the dura with a 14° angle) of 8–9-week-old wild-type mice with glass pipettes (tip diameter, 20–30 μm). Mice were then implanted bilaterally with custom-built optic fibers (diameter, 200 μm; numerical aperture, 0.39; Thorlabs) above the dmPFC (relative to the bregma, +2.1 mm AP, ±0.65 mm ML and −1.2 mm DV from the dura with a 14° angle). Before training in the aversive MIO paradigm, mice underwent a light-habituation session one day before the habituation session. During this light-habituation session, animals received 30 light stimulations of continuous green light (526.5 nm) during 7 s. Laser power at the source was set at 8–10 mW. Optogenetic tests were carried out with trained mice once performance reached stable levels (6–9 d of training). Light stimulation was applied for the duration of the CSs. The optogenetic test protocol included 30 CS^+^ light-on and 30 CS^+^ light-off intermingled and pseudorandomly ordered trials in which CS^+^1 and CS^+^2 were either paired (light-on) or not paired (light-off) with light stimulation. Likewise, there were 15 CS^−^ light-on and 15 CS^−^ light-off trials, intermingled with CS^+^. Two optogenetic test protocols were performed separated by a normal training session to control for the absence of long-term effects of optogenetic manipulation. For analysis, the number of transitions and percentage of time expressing Motif 2 were averaged across the two optogenetic test sessions for each mouse. Metrics quantified at the trial level, including transition probabilities, recurrence probabilities, motif density, and cumulative distributions, were calculated after pooling all trials across both optogenetic test sessions.

### Anatomical and histological analyses

Animals were euthanized under isoflurane anesthesia and perfused through the left ventricle with 4% (wt/vol) paraformaldehyde in 0.1 M PBS. Next, brains were dissected and post-fixed for 4 h at 4 °C in the same solution. Sections (60 μm thick) were cut, mounted on gelatin-coated microscope slides and dried. Sections were dehydrated and mounted. Electrolytic lesions were identified with conventional transmission light microscopy. Only recordings with confirmed lesions in cingulate and prelimbic areas of the dmPFC were included in our analyses. It is important to note that, because the electrode bundles are manually fabricated in the laboratory and individual wires can be slightly displaced during insertion in the brain, the distance between each electrode cannot be measured precisely. It is therefore technically impossible to determine from which layer the single units were recorded. For verification of viral infections in the dmPFC (cingulate and prelimbic areas), serial 80-μm-thick slices containing the dmPFC were imaged using an epifluorescence system (Leica DM5000) fitted with a ×10 dry objective. The location and the extent of the infections were visually assessed. Only infections accurately targeting the dmPFC were considered for behavioral and electrophysiological analyses.

### Surgeries for Ca^2+^ imaging

Eight-week-old mice were anesthetized using isoflurane (3 to 5% for induction, 1 to 2% for maintenance; Attane, Provet) in oxygen-enriched air (Oxymat 3, Weinmann) and placed in a stereotactic frame (model 1900, Kopf Instruments). Injections of buprenorphine (Temgesic, Indivior UK; 0.1 mg/kg body weight subcutaneously 30 min before anesthesia) and ropivacain (Naropin, AstraZeneca; 0.1 ml locally under the scalp before incision) were provided for analgesia. Postoperative pain medication included buprenorphine (0.1 mg/kg body weight in the drinking water overnight) and injections of meloxicam (Metacam, Boehringer Ingelheim; 1mg/kg subcutaneously) for 3 days after surgery. Ophthalmic gel was applied to avoid eye drying (Viscotears, Bausch + Lomb). The body temperature of the experimental animal was maintained at 36 °C using a feedback-controlled heating pad (FHC). AAV2/5.CaMK2.GCaMP6f (University of Pennsylvania Vector Core) was unilaterally injected into the dmPFC (300 nl) using precision micropositioner (model 2650, Kopf Instruments) and pulled glass pipettes (tip diameter ∼20 μm) connected to a Picospritzer III microinjection system (Parker Hannifin) at the following coordinates from bregma: anterior–posterior (AP): 2 mm; medial–lateral (ML): −0.5 mm; dorsal–ventral (DV): −1.4 mm. After injection, a track above the imaging site was cut with a sterile needle to aid the insertion of the GRIN lens (1 x 4 mm, Inscopix). The GRIN lens was subsequently lowered into the brain using a micropositioner through the track (4.3 mm from brain surface) with a custom-built lens holder and fixed to the skull using ultraviolet light-curable glue (Henkel, Loctite 4305). Dental acrylic (Paladur, Heraeus) was used to seal the skull and attach a custom-made head bar for animal fixation during the miniature microscope mounting procedure. Mice were allowed to recover for 3 weeks after surgery before checking for GCaMP expression.

### Ca^2+^ imaging recordings and processing

Mice were head-fixed to check GCaMP expression using a miniature microscope (nVista HD, Inscopix Inc.). If the expression level was sufficient, mice were anesthetized with isoflurane (3 to 5% for induction, 1 to 2% for maintenance; Attane, Provet) in oxygen-enriched air (Oxymat 3, Weinmann) to fix the miniature microscope baseplate (BLP-2, Inscopix) on top of the cranium implant using blue-light curable glue (Vertise Flow, Kerr). The miniature microscope was then detached, the baseplate was capped with a baseplate cover (Inscopix), and the mouse was returned to its home cage. During the 3 days before training, mice were habituated to head fixation (∼5 min), followed by a free exploration session in home cage (∼10 min). This daily procedure allowed mice to habituate to miniature microscope mounting and carrying. The miniature microscope was mounted immediately before each behavioral session by head fixation. Imaging data were acquired using nVista HD software (Inscopix Inc.) at a frame rate of 20 Hz (exposure time, 50 ms) with a light-emitting diode (LED) power of 0.6 to 0.8 mW/mm^2^ (excitation irradiance at objective front surface), an analog gain of 1 to 4, and a field of view of 1080 × 1080 pixels (∼648 × 648 μm with the nVista2 system) and 1280 × 800 pixels (∼768 × 480 μm, nVista3 system). Imaging frame timestamps were recorded with the OmniPlex neural recording data acquisition processor system.

Imaging files produced by the nVista2 or nVista3 systems were imported into MATLAB (Mathworks) using custom code. Two regions of interest (ROIs) manually selected by the investigators were used to correct for translational motion. To address background noise during motion correction, we subtracted a Gaussian-blurred image on a frame-by-frame basis from the raw imaging data. We used fast Fourier transform–based image registration^34^ to register the first 100 frames on one ROI, and then used the median image of these 100 frames as a template to register the rest of the frames in the imaging session until correction was lower than a user-defined threshold (<0.01 average pixel shift on average across all frames). The procedure was then repeated on the resulting motion-corrected movie using the second user-defined ROI to ensure minimal nonrigid motion. We applied calculated shifts to the raw movie and used the raw, motion-corrected movies for the extraction of Ca2+ traces and all subsequent analysis.

For trace extraction, we processed each session independently and used CNMF-E, an algorithm based on nonnegative matrix factorization^35^, for automatic extraction of Ca2+ traces. We then excluded from the analysis ROIs based on anatomy (ROI size, shape, or vicinity to the edge of lens), low signal-to-noise ratio or large overlap in signal and spatial location with other neurons (>60% spatial overlap and >0.6 Pearson’s correlation between the traces across the entire session). Linear trends across an entire session were removed from the Ca2+ traces and further calculations were performed on the *z* scores of the detrended traces.

### Single-unit analyses

Single-unit spike sorting was performed using Offline Sorter software (Plexon) and analyzed using NeuroExplorer (Nex Technologies) and MATLAB (Mathworks). Waveforms were manually defined while visualizing in a three-dimensional space using Principal Components (PCs), timing, and voltage features of the waveforms. A single unit was defined as a discrete, isolated cluster of waveforms in the feature space and did not contain spikes with a refractory period of less than 1 ms, based upon autocorrelation analyses. Additionally, multivariate ANOVA and J3 statistics were used to quantify separation of clusters in the PC space. Cross-correlation analyses were performed to control that a single unit was not recorded on multiple channels. Target neurons that displayed a peak of activity when the reference neuron fired were considered duplicates, and only a single neuron was considered for analysis. Single-unit instantaneous firing rate estimations were computed in 50 ms bins with a sliding window of 1 ms.

To investigate neuronal responses to the identified behavioral motifs, *z* scores were computed taking a reference period from −4 to 0 s before trial onset. For analysis of the percentage of neurons activated and inhibited by multiple behavioral motifs, we considered a neuron to be activated or inhibited if the *z* score was above 1.96 or below −1.96, respectively, for at least 250 ms. To further investigate the motif-response profiles of different neurons, we binned neuronal activity in time bins of 200 ms and considered a neuron to be active or inhibited in a time bin if its firing rate was above its basal firing rate (from −4 to 0 s before trial onset) plus three standard deviations or below its basal firing rate minus three standard deviations, respectively. A neuron was considered to be modulated by a motif if more than 70% of the samples were either activated or inhibited. We then classified each neuron into four categories according to their response profiles: selective (activated or inhibited by a single motif), mixed-motif responsive (activated by at least one motif and inhibited by at least one different motif), all responsive (activated or inhibited by all three motifs), and mixed modulated (mixture of activation and inhibitory responses to the same motif or motifs). For the identification of speed-modulated neurons, we computed the Pearson’s correlation coefficient r for each neuron and considered a neuron’s activity to be modulated by speed if the neuron was significantly correlated and its Pearson’s correlation coefficient r was above 0.1 or below −0.1.

To investigate the contribution of the recorded dmPFC neurons to behavioral motif transitions, we restricted our analysis to transition periods spanning at least 1.2 s. To detect the activation and inhibition times across behavioral transitions, we first computed the *z* score activity for each neuron taking a reference period from −4 to 0 s before trial onset. We then considered a neuron to be activated or inhibited if the *z* score was above 1.96 or below −1.96, respectively, for a minimum duration of 100 ms continuously. To compare the neuronal activity across behavioral transitions in success and failure trials, we subsampled trials to match the number of success and failure trials for each mouse before normalization, and used the average *z* score across 100 subsampling repetitions to perform the analysis described above. To compute the percentage of non-overlapping neurons between success and failure transition-modulated neurons, we calculated the percentage of neurons uniquely modulated during success trials (i.e., those not modulated during failure trials) over the total number of success-modulated neurons. We repeated the same analysis using two distinct subsets of success trials to compare these results to a within-condition control, providing an estimate of the degree of non-overlap expected by chance. The percentages of non-overlapping neurons were calculated separately for transitions from Motif 1 to Motif 2 and Motif 2 to Motif 3, and the global estimate was obtained by averaging across transition types.

### Population decoding

For population decoding analyses, firing activity from the ensemble of recorded neurons at a given time point *t* was pooled in a vector containing as many dimensions as cells recorded. The instantaneous pseudo-population vector (PV(t)) contains the activity of all the recorded neurons and mice at time *t*. The activity was binned in 200-ms time bins to match the sampling from the behavioral segmentation. To assess the decoding accuracies of the dmPFC population, we used linear-kernel support vector machine (SVM) classifiers. Single-trial instantaneous pseudo-population vectors were composed of the spike counts of each neuron using units and trials from different mice, therefore removing decoding contributions due to noise correlations. For the pairwise decoding of motif versus motif activity, the population vector at a random time *t* in relation to the expression of a behavioral motif was decoded from the population vector at a random time *t* of a different behavioral motif (Figure 2D). The decoding of motif versus pre-CS baseline activity was performed among population vectors for a given motif at a random time *t* in relation to population vectors during a baseline pre-CS period (Figure S3C). For the time-to-time decoding of behavioral transitions, we restricted our analysis to transition sequences of at least 1.2 s for the most frequent behavioral transitions, namely from Motif 1 to Motif 2 and Motif 2 to Motif 3 transitions. The population vector at time *t*n in relation to the onset of the behavioral transition was decoded from the population vector at the preceding time *t*n-1 (Figure 3C). For the decoding of behavioral transitions from motif activity, the population vector at time *t*n in relation to the onset of the behavioral transition was decoded from a population vector at a random time *t* of the motifs expressed during the transition (Figure S5B). Cross-temporal decoding of behavioral transitions was performed by doing the decoding procedure described above, using motif activity to decode behavioral transitions. Pseudo-population vectors were constructed in 100 ms bins aligned to the onset of behavioral transitions. Classifiers were trained to decode each transition time bin *t*n from a random time *t* during motif expression, and were tested with the same classes across all other transition time bins *t*n. This procedure was repeated for all combinations of train and test time bins in an intersectional manner, generating a cross-temporal decoding matrix in which each element represents decoding accuracy for a given train-test time-bin pair. This approach allowed us to determine how transition-related representations evolved over time. If representations are confined to a specific epoch of motif expression, decoding is expected only for the corresponding train-test time combinations, whereas decoding across additional train-test combinations indicates that motif-related information is represented outside that epoch. For the shifted cross-temporal decoding analysis, we used the same procedure but pseudo-population vectors were extracted from a window shifted 1 s preceding transition onset (−1.6 to −0.4 s) to perform cross-temporal decoding of population activity before the transition period (Figure S5G). For quantification of cross-temporal decoding, we computed weighted decoding accuracies for each test time bin column as the mean decoding accuracy across significantly decoded time bins in the test column, weighted by the proportion of significant bins. The proportion of significantly decoded test time bins was represented by horizontal shaded areas, with the height indicating the percentage of significant bins. For motif versus basal and motif versus motif decoding during ITI, we used the same procedure as described above but for motifs identified during an ITI period. To evaluate the generalization of classification accuracies to failure trials, we tested the classifiers with pseudo-population vectors from the distinct motifs expressed during failure trials (Figure S7A). To do this, we first trained a classifier to decode neuronal activity during success trials, and tested it using a dataset with the same classes from a failure trial, yielding the failure generalization decoding accuracy. To test the contribution of speed to the overall decoding of motif versus motif and motif versus pre-CS, we performed the same techniques as described above but removing speed-modulated neurons from the analysis.

To obtain comparable accuracies across the classifiers used in the different conditions, we used for all classifiers the same number of trials to train and test decoders. We used a cross-validation ‘holdout’ method that partitions the data into two mutually exclusive subsets, the training and test sets, with a 3:1 ratio. Using this method, we trained a classifier using the training dataset and evaluated the accuracy of the classifier in the test dataset. The decoding accuracy of a classifier was computed as the proportion of correctly classified trials in the test set. To estimate the variability of decoding accuracy, we performed 100 bootstrap iterations on randomly selecting trials for pseudo-population trial construction and randomly assigning them to the training and test dataset. The shuffled label decoding was obtained by randomly shuffling trial labels and used to generate a null distribution of the decoding accuracies that would occur by chance. This holdout cross-validation procedure was used for all decoding analyses, including those assessing the generalization of classification accuracy across success and failure trials. To assess statistical significance, trial labels were randomly shuffled and the decoding procedure was repeated 200 times. For each permutation, decoding accuracy was estimated as the average across 10 independent holdout iterations. For each time bin, P values were calculated as the proportion of shuffle repetitions that exceeded the real mean decoding accuracies, and significant mean decoding accuracies were defined as those time bins in which P<0.05.

### Population analyses

Time correlation analyses were performed separately for success and failure trials. For analyses performed across the entire behavioral sequence, population vectors were constructed separately for each mouse by combining the spiking activity of all recorded dmPFC units in 50-ms time bins. For each trial, a time-by-time Pearson correlation matrix was computed by correlating the population activity vectors across all pairs of time bins within the trial. Correlation matrices were restricted to behavioral sequence duration and resized to a common 200 x 200 grid to normalize for differences in trial duration. Correlation matrices were then averaged within each mouse to obtain a single time correlation matrix per mouse. Global correlation matrices were obtained by averaging matrices across mice. For quantification, global correlation matrices were restricted to a band centered on the main diagonal (± 75 resized bins). This region captured correlations where differences between conditions were most prominent, while minimizing contributions from correlations between temporally distant time points that could obscure the observed effects. The diagonal band was divided into five equally sized segments, and the mean correlation was computed by averaging all values contained within each segment. The resulting mean correlations were normalized using the combined values from success and failure trials, yielding normalized time correlation values on a common 0–1 scale for direct comparison between conditions.

To assess the relationship between dmPFC time correlation and Motif 2 expression, we fit a linear mixed-effects model using the percentage of time expressing Motif 2 as the dependent variable and dmPFC time correlation as the fixed-effects predictor. Each observation corresponded to a single trial. For each trial, dmPFC time correlation was quantified as the mean pairwise Pearson correlation between population activity vectors (containing the pooled activity of all recorded neurons within a mouse) across all time bins within the trial, excluding self-correlations. Time correlation values were mean-centered within each mouse before model fitting to remove between mouse differences in baseline time correlation and improve model interpretability. Models were fitted using the MATLAB function fitlme (Mathworks). Mouse identity was included as a random effect with random intercepts and random slopes to account for inter-mouse variability in both Motif 2 expression and the relationship between the predictor and dependent variable: Motif 2 expression ∼ time correlation + (time correlation | mouse).

To compute time correlation during matched behavioral transitions in success and failure trials, single-trial pseudo-population vectors were constructed by pooling the activity of all recorded dmPFC neurons across mice as described before for the decoding analyses. We restricted our analysis to transition sequences of at least 1.2 s. For each behavioral transition, time-by-time Pearson correlation matrices were computed by correlating population activity vectors across all pairs of time bins (50 ms), following the same procedure as described above. We repeated this process 100 times to obtain average correlation matrices. For quantification, we averaged the Pearson correlation coefficients across the entire correlation matrix and normalized the resulting values using the pooled success and failure trials, yielding a common 0–1 scale for comparison between conditions.

To quantify the separation between motif-related dmPFC activity, Mahalanobis distances were computed in the full dimensional PCA space. A pooled covariance matrix was estimated from all bootstrap pseudo-population vectors across motifs and outcomes, and its pseudoinverse was used to compute the Mahalanobis distance between each bootstrap pseudo-population vector and the centroid of the bootstrap distribution of the other motif. The reported distances correspond to the average between the distance from Motif 1 to the centroid of Motif 2 and the reciprocal distance from Motif 2 to the centroid of Motif 1.

### Dimensionality reduction

For dimensionality reduction, we used instantaneous pseudo-population vectors obtained from the *z* score activity (to avoid PCA being dominated by neurons with higher modulation depths^27^). For separation of behavioral motif-related activity in latent space, neuronal activity was binned into 200 ms intervals. Latent dimensions were then computed using activity from one time bin during the expression of each of the three behavioral motifs together with the three bins preceding CS onset, thereby capturing variability associated with both motif expression and pre-CS baseline activity. For dimensionality reduction of motif-related population activity across outcomes, we used the same structure as described above but with bins corresponding to success and failure trials. We then obtained these full-space pseudo-population vectors containing the average *z* score activity of each neuron in different time bins. Next, we used PCA in these datasets using each neuron as a dimension and each time bin as an observation. Using this technique, we obtained dimensions significantly explaining most of the variance and projected the pseudo-population vectors into these latent dimensions. To assess the robustness of our results and control for sampling variability, we employed a bootstrapping procedure in which, for 100 iterations, trials were randomly sampled with replacement from the original dataset. To ensure a common scaling across bootstrap iterations, the same mean and standard deviation, computed from all trials before resampling, were used to standardize every bootstrap pseudo-population vector. For PCA performed across outcomes, the number of trials per condition was fixed to the minimum number across conditions to avoid biases introduced by unequal sampling, yielding balanced pseudo-population vectors. These resampled bootstraps were projected into the different PC dimensions using the weights assigned to each neuron during the PCA performed on the complete dataset.

### Statistics and reproducibility

No statistical methods were used to predetermine sample size. Experiments were not randomized, and investigators were not blinded to allocation during experiments and outcome assessment. All behavioral experiments were controlled by computer systems, and data were collected in an automated and unsupervised manner.

We conducted all analyses using either custom routines written in MATLAB (Mathworks) or SigmaPlot (Systat Software). Figures were assembled using Adobe Illustrator (Adobe). For all datasets, normality was tested using the Kolmogorov–Smirnov test (a < 0.05) and homogeneity of variance with Levene’s test (a < 0.05) to determine whether parametric or nonparametric analyses were required. Parametric analyses included two-sided paired t-test, two-sided one-sample t-test, one-way ANOVA and repeated-measures one-way ANOVA with post hoc Tukey’s or Dunn’s test if a significant main effect or interaction was observed. For all tests, *P<0.05, **P<0.01, ***P<0.001 and not significant (NS) >0.5; significance was calculated after correction for multiple comparisons. When required, nonparametric Wilcoxon rank-sum, Wilcoxon signed-rank, two-sided permutation or Kolmogorov-Smirnov tests were used. To adjust for multiple hypothesis testing, we used Holm-Bonferroni correction. All data were collected in a computer-controlled, automated and unbiased manner.

