## Supplementary Figures for "Neuronal control of learned behavioral sequencing"

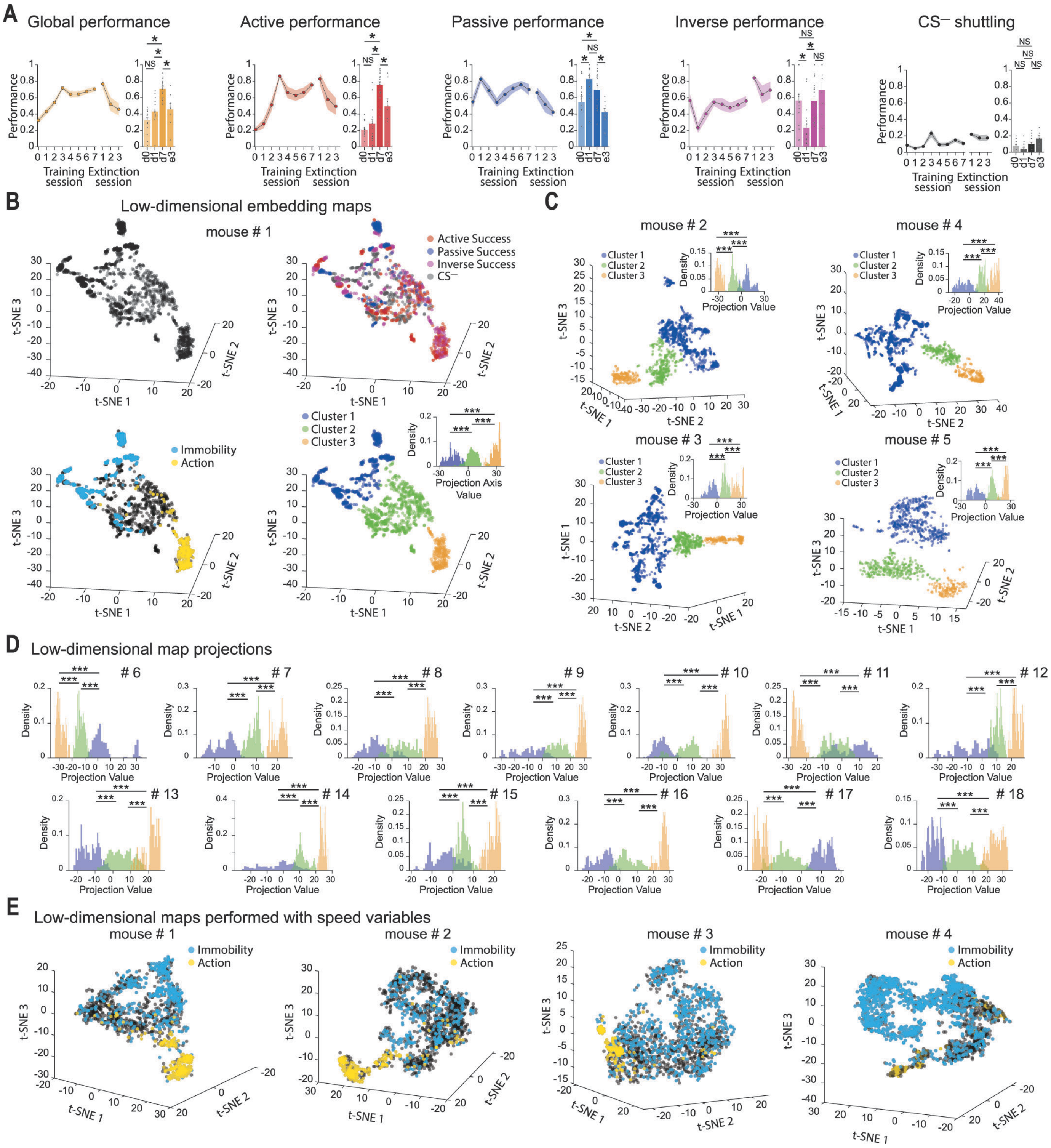

A
Global performance
Active performance
Passive performance
Inverse performance
CS⁻ shuttling
Performance
Training session
Extinction session
NS
B
Low-dimensional embedding maps
mouse # 1
Active Success
Passive Success
Inverse Success
CS⁻
Immobility
Action
Cluster 1
Cluster 2
Cluster 3
Projection Axis Value
Density
t-SNE 1
t-SNE 2
t-SNE 3
C
mouse # 2
mouse # 4
mouse # 3
mouse # 5
Projection Value
D
Low-dimensional map projections
# 6
# 7
# 8
# 9
# 10
# 11
# 12
# 13
# 14
# 15
# 16
# 17
# 18
E
Low-dimensional maps performed with speed variables
mouse # 1
mouse # 2
mouse # 3
mouse # 4

**Figure S1. Mouse performance across training and behavioral segmentation.** (**A**) Mean behavioral performance of n = 18 mice shown globally and for active, passive, inverse trials, and $CS^{-}$ shuttling, across training and extinction. Bar graphs show the comparison of mean performance during habituation (d0), the first day of training (d1), the last day of training (d7), and the last day of extinction (e3). Shaded areas and error bars represent S.E.M. One-way ANOVA with Tukey's or Dunn's post hoc test (*P<0.05; **P<0.01; ***P<0.001; NS=not significant). (**B**) Top left, low-dimensional embedding map of an example mouse. Top right, embedding with samples labeled by trial type (red: active success; blue: passive success; magenta: inverse success; black: control trials). Bottom left, embedding with samples labeled by the expression of immobility (light blue) and action (yellow). Bottom right, embedding with samples labeled by their assigned cluster (blue: Cluster 1; green: Cluster 2; orange: Cluster 3). Inset displays normalized sample density along the main projection axis for the identified clusters. (**C**) Low-dimensional embedding maps of four example mice. Embedding samples are labeled by their assigned cluster (blue: Cluster 1; green: Cluster 2; orange: Cluster 3). Insets display normalized sample density along the main projection axis for the identified clusters. Kolmogorov-Smirnov test (***P<0.001). (**D**) Main projection axis for the identified clusters of the remaining mice included in the analysis. Kolmogorov-Smirnov test (***P<0.001). (**E**) Low-dimensional embedding maps of four example mice generated exclusively from speed behavioral variables. Embedding samples are labeled by the expression of immobility (light blue) and action (yellow).

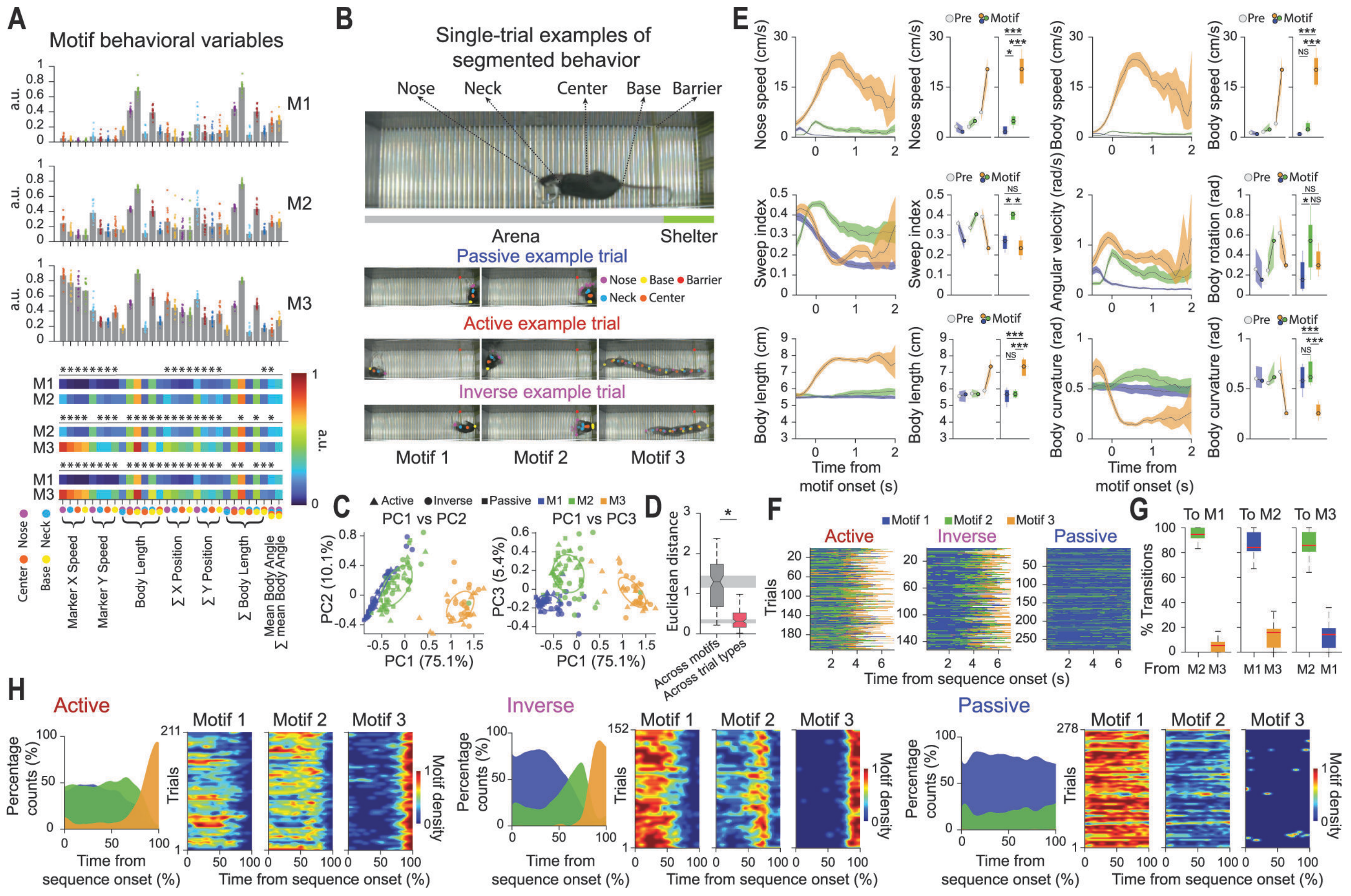
A
Motif behavioral variables
M1
M2
M3
a.u.
Nose
Neck
Center
Base
Marker X Speed
Marker Y Speed
Body Length
Σ X Position
Σ Y Position
Σ Body Length
Mean Body Angle
Σ mean Body Angle
B
Single-trial examples of segmented behavior
Nose
Neck
Center
Base
Barrier
Arena
Shelter
Passive example trial
Active example trial
Inverse example trial
Motif 1
Motif 2
Motif 3
C
Active
Inverse
Passive
PC1 vs PC2
PC1 vs PC3
PC2 (10.1%)
PC3 (5.4%)
PC1 (75.1%)
D
Euclidean distance
Across motifs
Across trial types
E
Nose speed (cm/s)
Body speed (cm/s)
Sweep index
Angular velocity (rad/s)
Body rotation (rad)
Body length (cm)
Body curvature (rad)
Pre
Motif
NS
Time from motif onset (s)
F
Motif 1
Motif 2
Motif 3
Active
Inverse
Passive
Trials
Time from sequence onset (s)
G
To M1
To M2
To M3
% Transitions
From
H
Active
Inverse
Passive
Percentage counts (%)
Trials
Motif density
Time from sequence onset (%)

**Figure S2. Behavioral motifs exhibit distinct behavioral signatures and are conserved across distinct trial types.** (**A**) Top, average levels of the behavioral variables used for behavioral segmentation for the different behavioral motifs (n = 18 mice). Bars represent mean, circles indicate individual mice. Bottom, comparison of behavioral variable levels across distinct behavioral motifs. Repeated measurements one-way ANOVA with post hoc Tukey and Holm-Bonferroni correction (*P<0.05). (**B**) Top, image of the behavioral box, divided into shelter and arena zones, with a mouse positioned inside. Bottom, example passive, active and inverse trials showing superimposed images and tracked body keypoints for each behavioral motif. (**C**) PCA projection of behavioral vectors across motifs and trial types in PC1–PC2 (left) and PC1–PC3 (right). Colors denote motifs (blue: Motif1; green: Motif 2; orange: Motif 3), and symbols denote trial types (triangle: active; circle: inverse; square: passive). (**D**) Euclidean distance between behavioral vectors in principal component space. Note that distances between behavioral motifs exceeded those between trial types within the same motif. Box plots show median (center line), 25th and 75th percentiles (box limits), 95% confidence intervals (notches), and 1.5xIQR (whiskers). Statistical significance was inferred from non-overlapping box-plot notches (*P<0.05). (**E**) Dynamics of behavioral variables aligned to motif onset (left) and corresponding quantification before and during motif expression (right). Repeated measurements one-way ANOVA with post hoc Tukey (*P<0.05; **P<0.01; ***P<0.001; NS=not significant). (**F**) Heatmaps of behavioral sequences across active, inverse and passive trials, aligned to sequence onset. (**G**) Percentage of transitions to each behavioral motif. Box plots show mean (center line), 25th and 75th percentiles (box limits), and 1.5xIQR (whiskers). (**H**) Behavioral sequences during active, inverse and passive trials. For each trial type, left panel shows the distribution of behavioral motifs across the sequences, showing the probability of observing each motif at each time point, and right panel shows corresponding motif-density heatmaps. Heatmaps were smoothed for visualization, and color intensity indicates motif density.

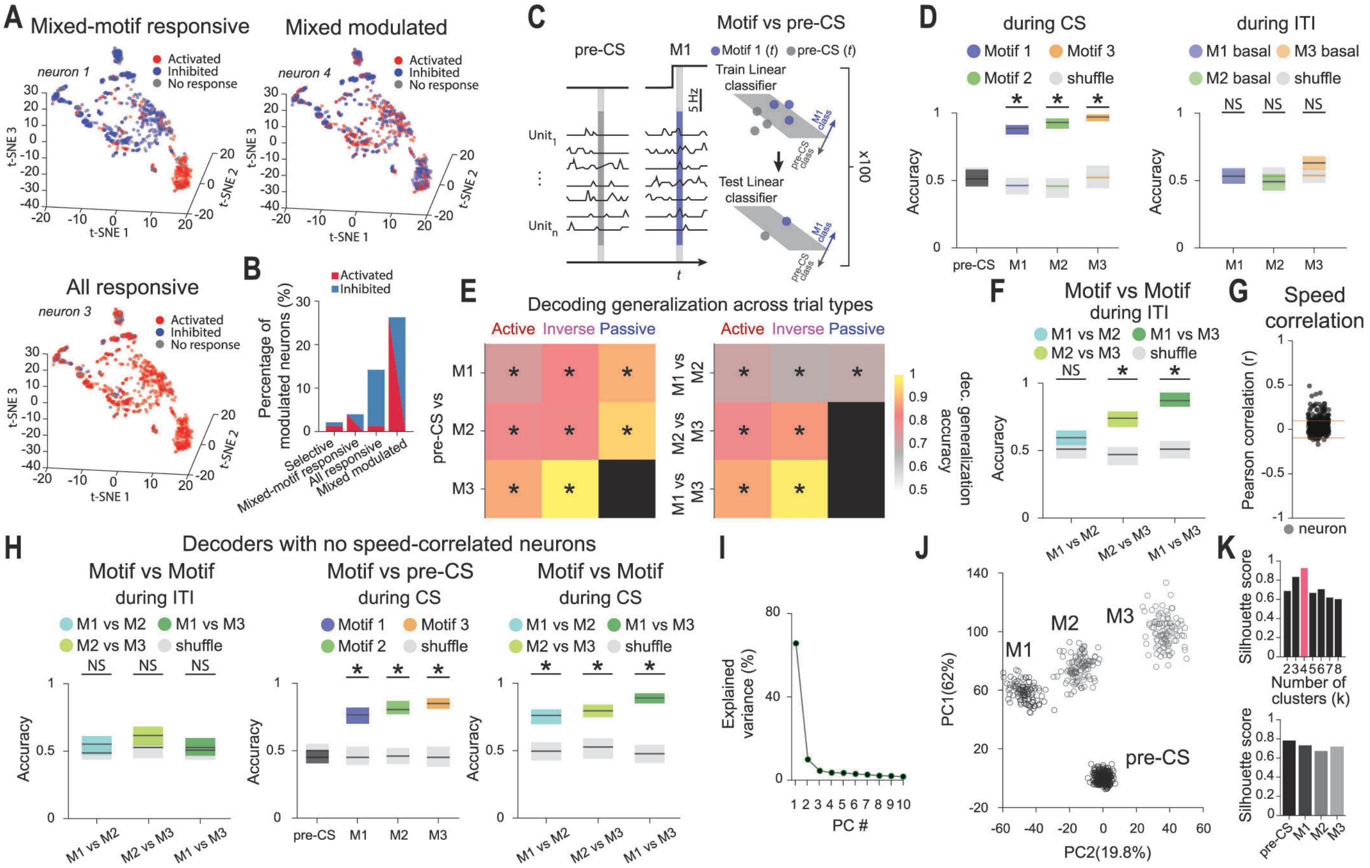
A
Mixed-motif responsive
Mixed modulated
All responsive
neuron 1
neuron 4
neuron 3
Activated
Inhibited
No response
t-SNE 1
t-SNE 2
t-SNE 3
B
Percentage of modulated neurons (%)
Selective
Mixed-motif responsive
All responsive
Mixed modulated
C
Motif vs pre-CS
pre-CS
M1
Motif 1 (t)
pre-CS (t)
Train Linear classifier
Test Linear classifier
M1 class
pre-CS class
x100
5 Hz
Unit1
Unitn
t
D
during CS
during ITI
Motif 1
Motif 3
Motif 2
shuffle
M1 basal
M3 basal
M2 basal
Accuracy
NS
E
Decoding generalization across trial types
Active
Inverse
Passive
pre-CS vs
M1 vs M2
M2 vs M3
M1 vs M3
dec. generalization accuracy
F
Motif vs Motif during ITI
M1 vs M2
M1 vs M3
M2 vs M3
G
Speed correlation
Pearson correlation (r)
neuron
H
Decoders with no speed-correlated neurons
Motif vs Motif during ITI
Motif vs pre-CS during CS
Motif vs Motif during CS
I
Explained variance (%)
PC #
J
PC1(62%)
PC2(19.8%)
K
Silhouette score
Number of clusters (k)

**Figure S3. Complementary analyses of behavioral motif representations.** (**A**) Single-neuron activity across the behavioral low-dimensional map (red: activated; blue: inhibited; gray: no response). Examples of mixed-motif responsive, all responsive and mixed modulated neurons. (**B**) Percentage of selective, mixed-motif responsive, all responsive and mixed modulated neurons. When applicable, neurons are subdivided into activated (red) and inhibited (blue). (**C**) Schematic of the decoding approach. The single trial-based pseudopopulation vector at time t is given by the spiking activity of the units for one trial. Linear classifiers are trained to classify motif from pre-CS baseline activity for trials at time *t* (Methods). (**D**) Mean decoding accuracies for classification of motifs from baseline activity during the CS period (left) and inter-trial intervals (right). Motif 1: M1, blue; Motif 2: M2, green; Motif 3: M3, orange; pre-CS: dark gray. (**E**) Cross trial-type decoding generalization of motif-related activity. Decoders trained on motif-related activity of two trial types were tested on a held-out trial type (active, inverse or passive trial), providing an estimate of the generalization of motif representations across distinct trial types. Left, motif versus pre-CS generalization accuracies. Right, motif versus motif generalization accuracies. (**F**) Motif versus motif decoding during inter-trial intervals. Decoding is shown for Motif 1 versus Motif 2 (blue), Motif 2 versus Motif 3 (light green) and Motif 1 versus Motif 3 (dark green). (**G**) Pearson correlation *r* between neuronal activity and speed. Circles represent individual neurons and red lines indicate thresholds ($r \pm 0.1$, *P<0.05). (**H**) Left, as in (F), but removing speed-correlated neurons. Middle, as in (D), left, but removing speed-correlated neurons. Right, as in (F), but for decoding during the CS period removing speed-correlated neurons. For (D, E, F and H), shuffle decoding accuracies (light gray) were obtained by randomizing the condition labels; shaded areas correspond to 95% confidence intervals; two-sided permutation test (*P<0.05). (**I**) Variance explained by principal components for the projection of motif activity in PCA space. (**J**) Projection of dmPFC population activity using PCA during pre-CS and motif-related activity in PC1 and PC2 dimensions. Circles represent resampled population vectors, and clusters were identified using k-means. (**K**) Top, silhouette scores for k-means clustering across different numbers of clusters *k*. Higher silhouette scores indicate greater separation and cohesion of the identified clusters, with the highest silhouette score observed for *k* = 4 (red). Bottom, silhouette scores for individual clusters using *k* = 4. For (D–G) and (I–K), n = 14 mice and 331 neurons. For (H), n = 14 mice and 248 neurons. Data shown are from success trials.

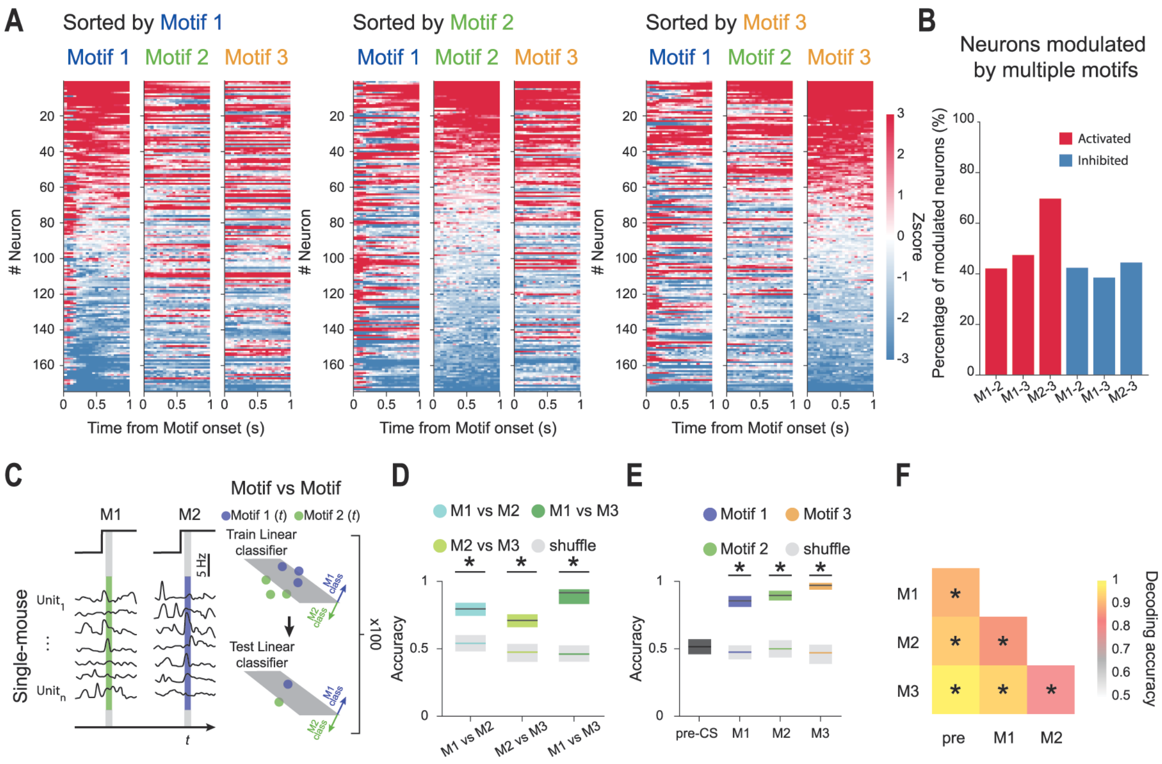

A
Sorted by Motif 1
Sorted by Motif 2
Sorted by Motif 3
Motif 1
Motif 2
Motif 3
### Neuron
Time from Motif onset (s)
Zscore
B
Neurons modulated by multiple motifs
Activated
Inhibited
Percentage of modulated neurons (%)
M1-2
M1-3
M2-3
C
Single-mouse
Unit1
Unitn
M1
M2
5 Hz
Motif vs Motif
Motif 1 (t)
Motif 2 (t)
Train Linear classifier
Test Linear classifier
M1 class
M2 class
x100
D
M1 vs M2
M1 vs M3
M2 vs M3
shuffle
Accuracy
E
Motif 1
Motif 2
Motif 3
shuffle
pre-CS
F
pre
M1
M2
M3
Decoding accuracy

**Figure S4. Single-mouse representations of discrete behavioral motifs.** (**A**) Heatmaps of *z* scored calcium activity aligned to motif onset (n = 174 neurons from 1 mouse). Neurons are sorted based on their activity at the onset of Motif 1 (left), Motif 2 (middle) and Motif 3 (right). Red and blue indicate increased and decreased activity, respectively. (**B**) Percentage of motif-modulated neurons responding to multiple motifs (red: activated; blue: inhibited). (**C**) Schematic of the decoding approach. The single trial-based pseudopopulation vector at time *t* is given by the calcium activity of the units recorded in a single-mouse for one trial. Linear classifiers are trained to classify Motif 1 and Motif 2-evoked population activity (Methods). (**D**) Mean decoding accuracies for pairwise classification of behavioral motifs based on dmPFC population activity. Decoding is shown for Motif 1 versus Motif 2 (blue), Motif 2 versus Motif 3 (light green) and Motif 1 versus Motif 3 (dark green). (**E**) Mean decoding accuracies for classification of behavioral motifs from pre-CS baseline activity. Decoding performance is shown for Motif 1 (blue), Motif 2 (green) and Motif 3 (orange), from pre-CS baseline periods (dark gray). For (D and E), shuffle decoding accuracies were obtained by randomizing the condition labels (light gray), and shaded areas correspond to 95% confidence intervals. (**F**) Summary of mean decoding accuracies for motif identity during the CS period. For (D–F), two-sided permutation test ($^*P<0.05$). Data shown are from success trials.

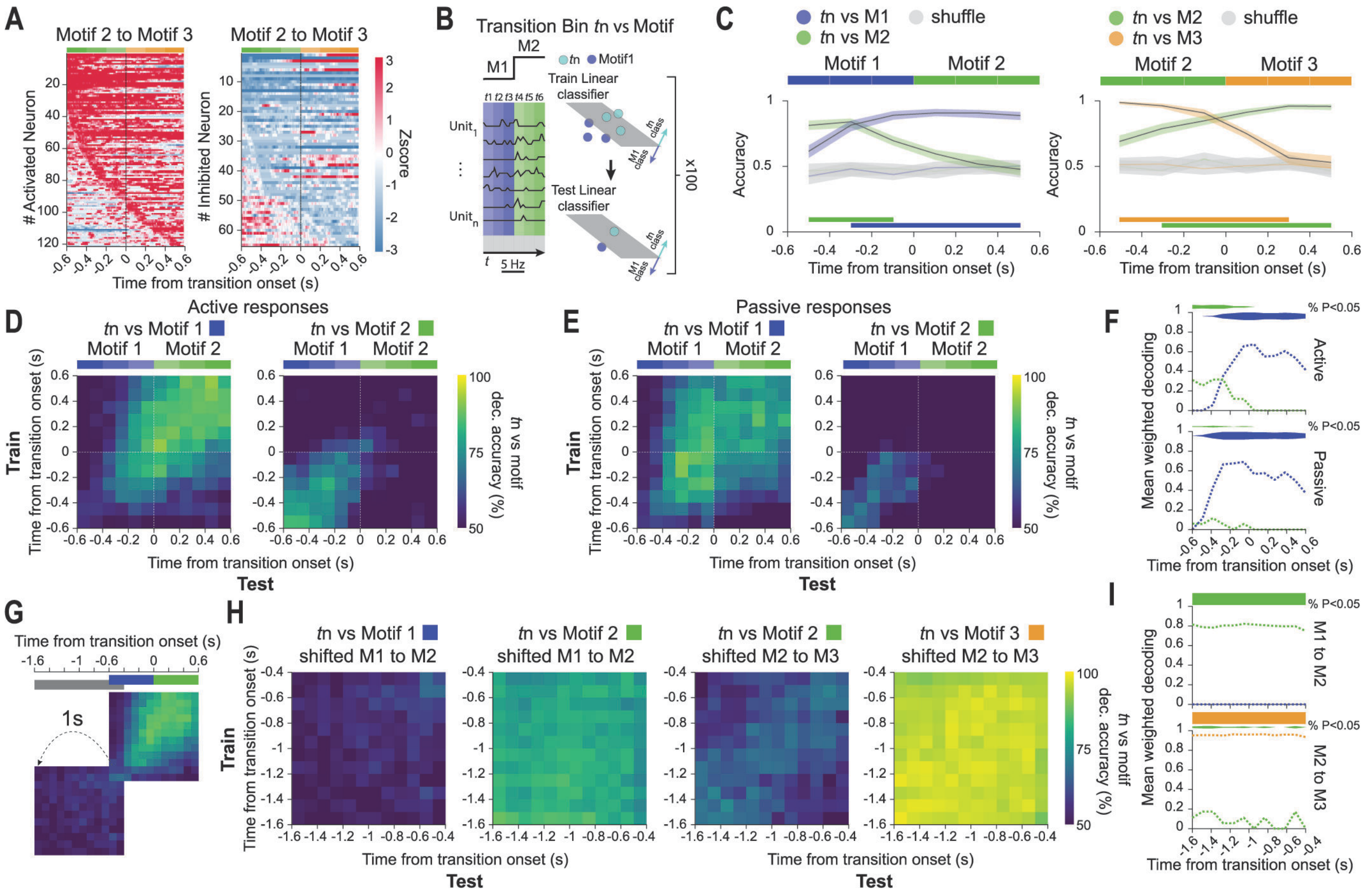

A
Motif 2 to Motif 3
Motif 2 to Motif 3
### Activated Neuron
### Inhibited Neuron
Zscore
Time from transition onset (s)
B
Transition Bin tn vs Motif
M1
M2
tn
Motif1
Train Linear classifier
Test Linear classifier
Unit1
Unitn
5 Hz
x100
C
tn vs M1
tn vs M2
shuffle
Motif 1
Motif 2
Accuracy
Time from transition onset (s)
tn vs M2
tn vs M3
shuffle
Motif 2
Motif 3
Accuracy
Time from transition onset (s)
D
Active responses
tn vs Motif 1
tn vs Motif 2
Motif 1
Motif 2
Train
Time from transition onset (s)
Test
tn vs motif dec. accuracy (%)
E
Passive responses
tn vs Motif 1
tn vs Motif 2
Motif 1
Motif 2
Train
Time from transition onset (s)
Test
tn vs motif dec. accuracy (%)
F
Mean weighted decoding
% P<0.05
Active
Passive
Time from transition onset (s)
G
Time from transition onset (s)
1s
H
tn vs Motif 1
shifted M1 to M2
tn vs Motif 2
shifted M1 to M2
tn vs Motif 2
shifted M2 to M3
tn vs Motif 3
shifted M2 to M3
Train
Time from transition onset (s)
Test
tn vs motif dec. accuracy (%)
I
Mean weighted decoding
% P<0.05
M1 to M2
M2 to M3
Time from transition onset (s)

**Figure S5. Complementary analyses of behavioral transition representations.** (**A**) Heatmaps of neuronal activity for activated (left) and inhibited (right) neurons, aligned to transitions from Motif 2 to Motif 3. Neurons are sorted by the first detected modulation time. Red and blue indicate increased and decreased activity, respectively. (**B**) Schematic of the decoding approach. The single-trial pseudopopulation vector at time *t* is given by the spiking activity of the units for one trial during a given behavioral transition. Linear classifiers are trained to classify each time bin *tn* (200-ms bin) of the transition from motif activity (Methods). (**C**) Mean decoding accuracies of transition time bins from motif activity during Motif 1 to Motif 2 (left) and Motif 2 to Motif 3 transitions (right). Shuffle decoding accuracies were obtained by randomizing the condition labels (light gray). Shaded areas correspond to 95% confidence intervals. Horizontal lines indicate significant decoding accuracies (two-sided permutation test; $P<0.05$). (**D**) Cross-temporal decoding of behavioral transitions (100-ms bin) from motif population activity for Motif 1 to Motif 2 transitions during active responses (active and inverse trials). Classifiers were trained at one time point and tested across all others for each transition bin (Methods). (**E**) As in (D), but during passive trials. (**F**) Quantification of cross-temporal decoding over time for transitions from Motif 1 to Motif 2 during active (top) and passive (bottom) response trials. Mean weighted decoding accuracy was calculated for each test time bin as the average decoding accuracy across significant bins in the corresponding column, weighted by the proportion of significant bins. Horizontal shaded areas indicate significant decoding, and height of shaded area indicates the percentage of decoded bins (two-sided permutation test; $P< 0.05$). (**G**) Schematic of shifted cross-temporal decoding using population activity 1 s before transition onset (-1.6 to -0.4 s). (**H**) As in (D), but for cross-temporal decoding using the shifted window for Motif 1 to Motif 2 (left) and Motif 2 to Motif 3 (right) transitions. (**I**) As in (F), but for the quantification of cross-temporal decoding using the shifted window for Motif 1 to Motif 2 (top) and Motif 2 to Motif 3 (bottom) transitions. Two-sided permutation test ($P<0.05$). Data shown are from success trials. n = 318 neurons from 13 mice.

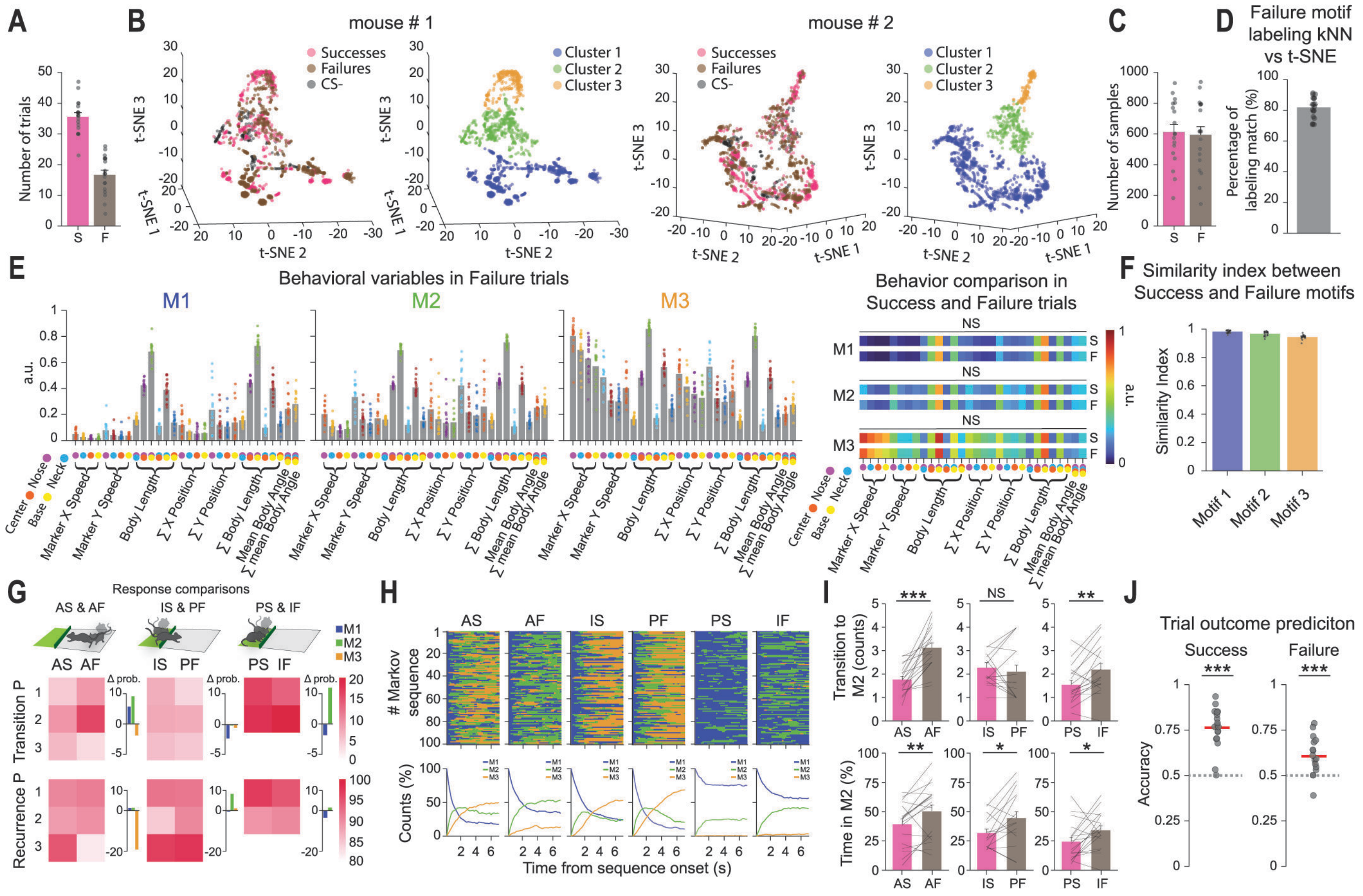
A
B
mouse # 1
mouse # 2
Successes
Failures
CS-
Cluster 1
Cluster 2
Cluster 3
t-SNE 1
t-SNE 2
t-SNE 3
Number of trials
S
F
C
Number of samples
D
Failure motif labeling kNN vs t-SNE
Percentage of labeling match (%)
E
Behavioral variables in Failure trials
M1
M2
M3
a.u.
Center
Nose
Base
Neck
Marker X Speed
Marker Y Speed
Body Length
∑ X Position
∑ Y Position
∑ Body Length
Mean Body Angle
∑ mean Body Angle
Behavior comparison in Success and Failure trials
NS
F
Similarity index between Success and Failure motifs
Similarity Index
Motif 1
Motif 2
Motif 3
G
Response comparisons
AS & AF
IS & PF
PS & IF
Transition P
Recurrence P
Δ prob.
H
AS
AF
IS
PF
PS
IF
### Markov sequence
Counts (%)
Time from sequence onset (s)
I
Transition to M2 (counts)
Time in M2 (%)
J
Trial outcome prediciton
Success
Failure
Accuracy

**Figure S6. Conserved behavioral motifs across outcomes and failure-specific sequence organization across trial types.** (**A**) Number of success (pink) and failure (brown) trials. Bars represent mean ± S.E.M., circles indicate individual mice. (**B**) Low-dimensional embedding maps performed with all success and failure trials are shown for two example mice, labeled by trial type (left, pink: success; brown: failure; black: $CS^{-}$), and by assigned cluster (blue: Cluster 1; green: Cluster 2; orange: Cluster 3). (**C**) Number of samples after downsampling to balance success and failure trials before segmentation. (**D**) Percentage of samples assigned to the same motif by k-NN interpolation or t-SNE embedding (Methods). (**E**) Left, average levels of the behavioral variables for the different behavioral motifs during failure trials. Right, comparison between success and failure trials. Paired t-test with Holm-Bonferroni correction (NS=not significant). (**F**) Similarity index comparing the behavioral profiles of success and failure trials across motifs. Similarity was quantified as one minus the mean absolute difference across all behavioral variables between success and failure trials. (**G**) Transition and recurrence probabilities across distinct trial types. Top, response-coupled trial comparisons (active success versus active failure; inverse success versus passive failure; passive success versus inverse failure). Bottom, transition and recurrence probabilities across response-coupled success and failure trials and their differences. (**H**) Top, heatmaps of surrogate Markov sequences generated from the HMM-estimated transition probability matrix. Bottom, percentage of motif occurrences over time for the different trial types. (**I**) Number of transitions to Motif 2 (top) and time spent in Motif 2 (bottom) across response-coupled success and failure trials. Bars represent mean ± S.E.M., lines indicate individual mice. Paired t-test (*$P<0.05$; **$P<0.01$; ***$P<0.001$; NS=not significant). (**J**) Trial outcome prediction from behavioral sequences of success (left) and failure (right) trials. Circles indicate held-out mice and red line indicates mean accuracy across mice. One-sample t-test against chance (0.5; ***$P<0.001$); n = 18 mice.

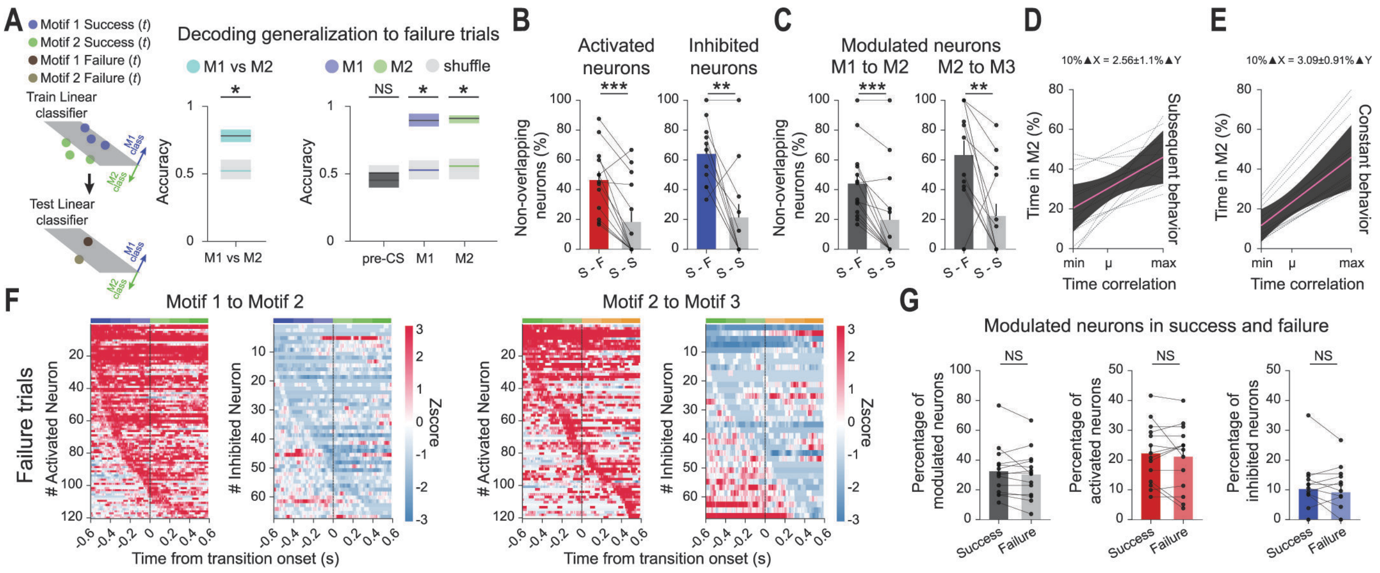
A
Motif 1 Success (t)
Motif 2 Success (t)
Motif 1 Failure (t)
Motif 2 Failure (t)
Train Linear classifier
Test Linear classifier
Decoding generalization to failure trials
M1 vs M2
M1 M2 shuffle
Accuracy
NS
pre-CS
M1
M2
B
Activated neurons
Inhibited neurons
Non-overlapping neurons (%)
S - F
S - S
C
Modulated neurons
M1 to M2
M2 to M3
D
10%▲X = 2.56±1.1%▲Y
Time in M2 (%)
Subsequent behavior
Time correlation
E
10%▲X = 3.09±0.91%▲Y
Constant behavior
F
Motif 1 to Motif 2
Motif 2 to Motif 3
Failure trials
### Activated Neuron
### Inhibited Neuron
Zscore
Time from transition onset (s)
G
Modulated neurons in success and failure
Percentage of modulated neurons
Percentage of activated neurons
Percentage of inhibited neurons
Success
Failure
NS

**Figure S7. Complementary analyses of outcome-dependent dmPFC activity patterns during behavioral transitions.** (**A**) Left, decoding generalization strategy. Linear classifiers are trained to classify Motif 1 versus Motif 2-evoked population activity during success trials and tested during failure trials (Methods). Middle, mean generalization decoding accuracies for classification of Motif 1 versus Motif 2. Right, mean generalization decoding accuracies for classification of motif activity from pre-CS baseline activity. Shuffle decoding accuracies (light gray) were obtained by randomizing the condition labels. Shaded areas correspond to 95% confidence intervals. Two-sided permutation test (*$P<0.05$). (**B**) Percentage of non-overlapping transition-modulated neurons across outcomes versus a within-condition control, subdivided into activated (left) and inhibited (right) neurons. Non-overlapping neurons were modulated in one condition but not the other. Bars represent mean, circles individual mice. (**C**) As in (B), but subdivided in transitions from Motif 1 to Motif 2 (left), and Motif 2 to Motif 3 (right). (**D**) Relationship between dmPFC time correlation (mean-centered; $\mu$) and subsequent Motif 2 expression during success trials estimated with a linear mixed-effects model (Methods). Black lines indicate individual mouse fits, pink line indicates the population estimate, and shaded area represents the 95% confidence interval. A 10% increase in time correlation predicted a 2.56 ± 1.1% increase in subsequent Motif 2 expression (LME: $\beta = 35.2 \pm 15.2$, $t = 2.32$, $P = 0.02$). (**E**) As in (D), but matching behavior across trials. A 10% increase in time correlation predicted a 3.09 ± 0.91% increase in Motif 2 expression (LME: $\beta = 32 \pm 9.4$, $t = 3.41$, $P<0.001$). (**F**) Heatmaps of neuronal activity for activated (red) and inhibited (blue) neurons during failure trial aligned to transitions from Motif 1 to Motif 2 (left) and Motif 2 to Motif 3 (right). Neurons are sorted by the first detected modulation time. Red and blue indicate increased and decreased activity, respectively. (**G**) Percentage of modulated, activated, and inhibited neurons during behavioral transitions in success and failure trials. For (B, C and G), paired t-test or Wilcoxon signed-rank test (**P<0.01; ***P<0.001; NS=not significant). For (A–G), n = 331 neurons from 14 mice.

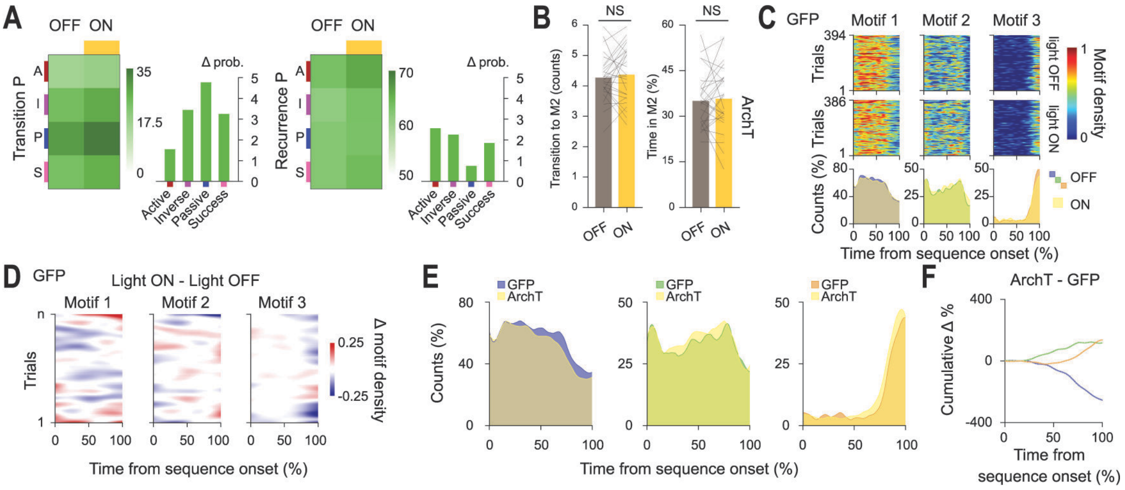
A
OFF ON
Transition P
Δ prob.
Active
Inverse
Passive
Success
Recurrence P
B
NS
Transition to M2 (counts)
Time in M2 (%)
OFF ON
ArchT
C
GFP
Motif 1
Motif 2
Motif 3
Trials
light OFF
light ON
Motif density
Counts (%)
OFF
ON
Time from sequence onset (%)
D
GFP
Light ON - Light OFF
Δ motif density
E
GFP
ArchT
Counts (%)
Time from sequence onset (%)
F
ArchT - GFP
Cumulative Δ %
Time from sequence onset (%)

**Figure S8. Complementary analyses of dmPFC optogenetic inhibition.** (**A**) Transition (left) and recurrence (right) probabilities for Motif 2 across light-off and light-on conditions in active (A, red), inverse (I, magenta), passive (P, blue) and overall (S, pink) trials, and quantification of the differences between stimulation conditions. (**B**) Left, number of transitions to Motif 2 during failure light-off (brown) and light-on (yellow) trials in the ArchT group. Right, percentage of time expressing Motif 2 during failure light-off and light-on trials in the ArchT group. Bars represent mean ± S.E.M., lines indicate individual mice. Paired t-test (NS=not significant). (**C**) Top, motif-density heatmaps showing the occurrence of behavioral motifs across behavioral sequences in light-off and light-on conditions in the GFP control group. Color intensity indicates motif density. Matrices were smoothed for visualization. Bottom, distribution of behavioral motifs over time in light-off (blue: Motif 1; green: Motif 2; orange: Motif 3) and light-on (yellow) conditions in the GFP control group. (**D**) Difference heatmaps between light-on and light-off conditions in the GFP control group. Red and blue indicate increased and decreased motif density during light delivery, respectively. (**E**) Distribution of behavioral motifs over behavioral sequences in the GFP control (blue: Motif 1; green: Motif 2; orange: Motif 3) and ArchT (yellow) groups during light-on trials. (**F**) Cumulative difference in motif probability between ArchT and GFP control groups during light-on trials. n = 29 ArchT mice and 18 GFP mice.

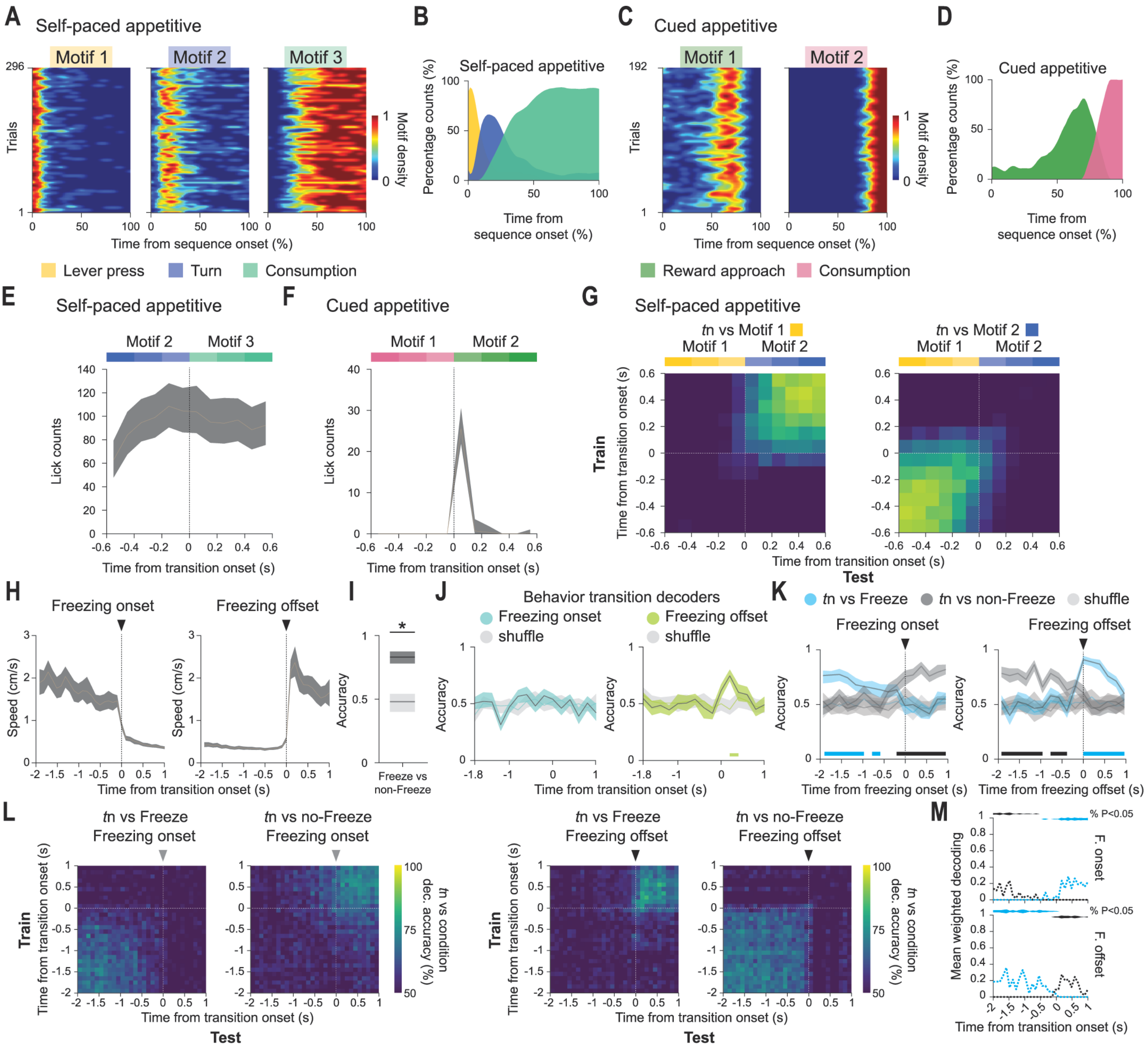
A Self-paced appetitive
Motif 1
Motif 2
Motif 3
Trials
Motif density
Time from sequence onset (%)
B Self-paced appetitive
Percentage counts (%)
Time from sequence onset (%)
C Cued appetitive
Motif 1
Motif 2
Trials
Motif density
Time from sequence onset (%)
D Cued appetitive
Percentage counts (%)
Time from sequence onset (%)
Lever press
Turn
Consumption
Reward approach
Consumption
E Self-paced appetitive
Motif 2
Motif 3
Lick counts
Time from transition onset (s)
F Cued appetitive
Motif 1
Motif 2
Lick counts
Time from transition onset (s)
G Self-paced appetitive
tn vs Motif 1
tn vs Motif 2
Train
Time from transition onset (s)
Test
H Freezing onset
Freezing offset
Speed (cm/s)
Time from transition onset (s)
I Accuracy
Freeze vs non-Freeze
J Behavior transition decoders
Freezing onset
Freezing offset
shuffle
Accuracy
Time from transition onset (s)
K tn vs Freeze
tn vs non-Freeze
shuffle
Freezing onset
Freezing offset
Accuracy
Time from freezing onset (s)
Time from freezing offset (s)
L tn vs Freeze Freezing onset
tn vs no-Freeze Freezing onset
tn vs Freeze Freezing offset
tn vs no-Freeze Freezing offset
Train
Time from transition onset (s)
Test
tn vs condition dec. accuracy (%)
M Mean weighted decoding
% P<0.05
F. onset
F. offset
Time from transition onset (s)

**Figure S9. Behavioral sequence structure during appetitive tasks and neuronal representations of freezing transitions.** (**A**) Motif density heatmaps showing the occurrence of behavioral motifs across behavioral sequences in a self-paced appetitive task. Motif 1, lever press; Motif 2, turn; Motif 3, consumption. Color intensity indicates motif density. Matrices were smoothed for visualization. (**B**) Distribution of behavioral motifs across the behavioral sequences, showing the probability of observing each motif at each time point. (**C** and **D**), As in (A and B), but for behavioral sequences in a cued appetitive task. Motif 1, reward approach; Motif 2, consumption. (**E**) Instantaneous lick count sum aligned to transitions from turn (Motif 2) to consumption (Motif 3) in a self-paced appetitive task. (**F**) As in (E), but for transitions from reward approach (Motif 1) to consumption (Motif 2), in a cued appetitive task. (**G**) Cross-temporal decoding (100-ms bin) for transitions from Motif 1 to Motif 2 in a self-paced appetitive task. Classifiers were trained at one time point and tested across all others for each transition bin (Methods). (**H**) Median instantaneous speed aligned to freezing onset (left) and freezing offset (right), during the retrieval session after contextual fear conditioning. (**I**) Mean decoding accuracy for classification of freezing versus non-freezing periods. Two-sided permutation test ($^*P<0.05$). (**J**) Decoding accuracies across consecutive time bins (200-ms bin) during freezing onset (left) and freezing offset (right) transitions. (**K**) Mean decoding accuracies obtained for the classification of transition time bins from freeze (blue) and non-freeze (black) activity periods, aligned to freezing onset (left) and offset (right). For (J and K), shuffle decoding was obtained by randomizing condition labels (light gray); shaded areas correspond to 95% confidence intervals; horizontal lines indicate significant decoding (two-sided permutation test; $P< 0.05$). (**L**) Cross-temporal decoding of freezing transitions (100-ms bin) from freeze (blue) and non-freeze (black) activity periods. Data are aligned to freezing onset (left) and freezing offset (right). (**M**) Quantification of cross-temporal decoding for freezing onset (top) and freezing offset (bottom) transitions. Mean weighted decoding was calculated from significant time bins and weighted by the proportion of significantly decoded bin. Horizontal shaded areas indicate significant decoding, and height of shaded area indicates the percentage of decoded bins (two-sided permutation test; $P < 0.05$). For (A, B and E), n = 10 mice. For (C, D and F), n = 4 mice. For (G), n = 10 mice and 1776 neurons. For (H–M), n = 19 mice and 409 neurons.
